# Beyond rates: Time-varying dynamics of high frequency oscillations as a biomarker of the seizure onset zone

**DOI:** 10.1101/2020.05.28.122416

**Authors:** Michael D. Nunez, Krit Charupanit, Indranil Sen-Gupta, Beth A. Lopour, Jack J. Lin

## Abstract

**Objective:** High frequency oscillations (HFOs) recorded by intracranial electrodes have generated excitement for their potential to help localize epileptic tissue for surgical resection. However, the number of HFOs per minute (i.e. the HFO “rate”) is not stable over the duration of intracranial recordings; for example, the rate of HFOs increases during periods of slow-wave sleep. Moreover, HFOs that are predictive of epileptic tissue may occur in oscillatory patterns due to phase coupling with lower frequencies. Therefore, we sought to further characterize between-seizure (i.e. “interictal”) HFO dynamics both within and outside the seizure onset zone (SOZ).

**Approach:** Using long-term intracranial EEG (mean duration 10.3 hours) from 16 patients, we automatically detected HFOs using a new algorithm. We then fit a hierarchical Negative Binomial model to the HFO counts. To account for differences in HFO dynamics and rates between sleep and wakefulness, we also fit a mixture model to the same data that included the ability to switch between two discrete brain states that were automatically determined during the fitting process. The ability to predict the SOZ by model parameters describing HFO dynamics (i.e. clumping coefficients and coefficients of variation) was assessed using receiver operating characteristic curves.

**Main results:** Parameters that described HFO dynamics were predictive of SOZ. In fact, these parameters were found to be more consistently predictive than HFO rate. Using concurrent scalp EEG in two patients, we show that the model-found brain states corresponded to (1) non-REM sleep and (2) awake and rapid eye movement sleep. However the brain state most likely corresponding to slow-wave sleep in the second model improved SOZ prediction compared to the first model for only some patients.

**Significance:** This work suggests that delineation of seizure onset zone with interictal data can be improved by the inclusion of time-varying HFO dynamics.

**Novelty & Significance:** The rate of high frequency oscillations (HFOs), measured as number per minute, is a biomarker of the seizure onset zone (SOZ) in epilepsy patients. However, the rate changes over time and HFO occurrence can be phase-coupled to slow oscillations. Here we show, through novel application of negative binomial models to HFO count data, that HFO temporal dynamics are a biomarker of the SOZ and are superior to HFO rate. Specifically, more random occurrence of HFOs predicted SOZ, as opposed to events clustered in time. This suggests that consideration of HFO temporal dynamics can improve SOZ localization for epilepsy surgery.

## 2. Introduction

Epilepsy is prevalent across the globe. For example, 1.2% of the population of the United States in 2015 were reported to have epilepsy (Zack and Kobau, 2017). Of this multitude, about 30% to 40% have seizures that cannot be controlled by antiseizure medication (Kwan and Brodie, 2000; Engel, 2018). In such cases of drug-resistant epilepsy, seizures can greatly decrease the patient’s quality of life. However, surgical interventions such as resection of seizure-generating tissue and implantation of responsive neurostimulators (RNS) are procedures that can greatly reduce or eradicate the occurrence of seizures (Engel, 2018). The goal of epilepsy surgery is to identify and treat the epileptogenic zone (EZ), typically defined as the minimum amount of tissue that must be surgically removed or stimulated to achieve a seizure free outcome (Rosenow and Lüders, 2001; Kovac et al., 2017). However, the EZ is a theoretical construct, and no biomarkers exist that can accurately and consistently identify the EZ (Ryvlin et al., 2014). One method to approximate the EZ is to use intracranial electroencephalography (iEEG) to localize the seizure onset zone (SOZ) (Kovac et al., 2017), and the SOZ is then used in conjunction with other imaging and test results to select brain tissue for treatment (e.g. Tomás et al., 2019). While surgery often results in a reduction of seizures, many patients will not be seizure free, indicating that there is a need for more accurate methods of identifying the EZ (Noachtar and Borggraefe, 2009). Such improvements would allow more patients to benefit from this procedure, with fewer side effects from the surgery and better outcomes (especially those with epilepsy outside of the temporal lobe with normal MRIs; Cohen-Gadol et al., 2006; Noe et al., 2013).

High frequency oscillations (HFOs) have shown promise as a novel marker of the EZ. Specifically, increased incidence (i.e. increased “rate” per minute) of transitory HFOs (Bragin et al., 1999, 2002) is thought to be indicative of the EZ (Jacobs et al., 2008, 2010; Frauscher et al., 2017). HFOs are “transitory,” as they are defined as temporally isolated events that last less than 200 ms with 3 or more cycles (i.e. 6 positive and negative local peaks in the waveform; Staba et al., 2002; Jacobs et al., 2008; Charupanit and Lopour, 2017). HFOs are often subcategorized as ripple band (80 - 250 Hz) and fast ripple band (250 - 500 Hz) events, and unsupervised analysis of high frequency data has produced evidence for these two HFO subtypes (see Blanco et al., 2010). These waveforms are thought to be generated by synchronous population firing and/or synchronous postsynaptic activity in the brain, although there is an abundance of possible neural mechanisms and cortical circuits that could generate HFOs (Köhling and Staley, 2011; Staba and Bragin, 2011; Jefferys et al., 2012).

Research has further sought to differentiate *pathological* HFOs, occurring in the EZ, from *physiological* HFOs, which can occur across the brain due to normal neural processes. The difficulty in differentiating pathological HFOs from normal brain activity has been a barrier to the use of HFOs in modern clinical practice (Jacobs et al., 2018; Fedele et al., 2019). For example, even though high rates of HFOs are typically thought to be indicative of the SOZ, baseline rates of HFOs outside the SOZ vary across different regions of the cortex (Guragain et al., 2018; Frauscher et al., 2018). Pathological and physiological HFOs are also affected by the sleep state of the patient, and HFO rates during slow wave sleep (i.e. non-rapid eye movement; NREM sleep) are thought to be more differentiating of pathological versus physiological brain activity (Dümpelmann et al., 2015; von Ellenrieder et al., 2016, 2017). Fast ripples are generally more localized to SOZ than ripples (although see Jacobs et al., 2018; King-Stephens, 2019), but they occur less frequently and may not be recorded in all patients (Köhling and Staley, 2011; Roehri et al., 2018). Roehri et al. (2018) show that HFOs are not more predictive of SOZ than pathological epileptiform discharges, although the co-occurrence of both is most predictive. Gliske et al. (2018) found that ripples during NREM sleep are only predictive of SOZ in some patients, and that HFO sources were highly variable over time. This led Gliske et al. (2018) to make the argument that long recordings over multiple days must be performed in order to accurately measure interictal, ripple-band HFO dynamics.

Analyses of phase-amplitude coupling in iEEG suggest that the temporal dynamics of HFOs and the precise timing of their occurrence may be an important marker of epileptogenic tissue. Coupling of ripple-band HFOs to slow waves has been observed during preictal and seizure periods (Weiss et al., 2013; Ibrahim et al., 2014; Guirgis et al., 2015). Moreover, pathological, interictal HFOs may be modulated by high amplitude, low frequency background activity, especially during sleep (Kerber et al., 2014; Frauscher et al., 2015; von Ellenrieder et al., 2016; Song et al., 2017; Motoi et al., 2018). However, this characteristic of high frequency activity remains relatively unexplored compared to the simple counting of HFOs per minute.

In this study, we show that the temporal dynamics of HFOs, beyond the changing of HFO rate with sleep stage, are predictive of SOZ. In particular, the more Poisson-like the HFO generator, the more likely that tissue is to be in the SOZ as judged by area under the curves (AUCs) of receiver operating characteristic curves (ROCs). Tissue that generates HFOs occurring close together in time with long intermediate periods (e.g. temporal “clumping” of HFOs) is less likely to be part of the SOZ. We found this to be true in general across many hours of iEEG in 16 patients as well as in empirically-found brain states that are reflective of NREM sleep in those patients.

## 3. Materials & Methods

### 3.1 Ethics statement

Approval for this study was obtained from the Institutional Review Board of the University of California, Irvine.

### 3.2 Patients and iEEG recordings

Patients who had intractable epilepsy and were candidates for resective surgery had intracranial electrodes implanted at the University of California Irvine Medical Center to aid SOZ localization. We analyzed iEEG data from patients (*N* = 16, 8 female, 36 ± 15 years of age, see **Table 1**) who were implanted with either subdural electrocorticography (ECoG) grids or strips, depth macroelectrodes and/or stereotactic EEG (SEEG). The electrode types and locations were chosen by the clinicians for diagnostic and surgical evaluation.

**Table 1:** Clinical information for each patient including: age, gender (G), epilepsy diagnosis, surgery performed, Engel outcome (Engel, 1993), electrode types (Elec), number of SOZ channels used in model fitting (SOZ), number of non-SOZ channels used in model fitting (nSOZ), consecutive interictal hours of iEEG used in the model fitting, and the start time of the consecutive hours used in the model fitting in 24-hour format. * denotes patients for whom all channels were used to fit data to the mixture models, while we included only grey matter localized channels for the other patients. Abbreviations: P = partial, L = left, R = right, B = bilateral, F = frontal, T = temporal, L = lobe, E = epilepsy, S = sclerosis, CD = cortical dysplasia, A = reactive astrocytosis and cell loss, Lo = lobectomy, Le = lesionectomy, *RNS* = implanted responsive neurostimulator. Patients 5 and 6 had unknown Engel Outcomes. The electrode types were: D = depth including SEEG, S = subdural electrocorticography (ECoG) grids or strips, M = a mix of both types. ^1^Confirmed using surgical pathology in these patients. ^2^This patient also had hypothalamic hamartoma ablation performed. ^2^These patients had delta power that was significantly different with *p <* .001 between the two states in Model 2 using a Kruskal-Wallis test. ^$^ and ^#^ indicate the same significance information when using cutoffs *α* = .01 and *α*= .05 respectively.

| Patient | Age | G | Diagnosis | Surgery | Engel | Elec | SOZ | nSOZ | Hours | Start |
| --- | --- | --- | --- | --- | --- | --- | --- | --- | --- | --- |
| **1** | 50 | M | RTLE | RTL<sub>o</sub> | IB | D | 4 | 41 | 13 | 21:15<sup>+</sup> |
| **2** | 23 | F | LTLE | LTL<sub>o</sub> | IIIA | D | 4 | 44 | 25 | 15:40<sup>+</sup> |
| **3** | 34 | M | RTLE, BTLS | RTL<sub>o</sub> | IA | D | 3 | 41 | 15 | 18:05 |
| **4** | 21 | M | RTLE, A<sup>1</sup> | RTL<sub>o</sub><sup>2</sup> | IIB | M | 13 | 110 | 16 | 16:23<sup>§</sup> |
| **5\*** | 21 | F | RFLE | *RNS* | ? | S | 9 | 111 | 12 | 16:51<sup>+</sup> |
| **6\*** | 28 | F | RFLE, CD<sup>1</sup> | RFL<sub>e</sub>, PRFL<sub>o</sub> | ? | M | 4 | 92 | 5 | 17:08 |
| **7\*** | 28 | M | LTLE | LTL<sub>o</sub> | IIIA | D | 13 | 113 | 9 | 09:53 |
| **8\*** | 44 | M | RFLE, CD<sup>1</sup> | RFL<sub>o</sub> | IA | S | 5 | 123 | 12 | 17:19 |
| **9** | 57 | F | LTLE | LTL<sub>o</sub> | IA | D | 5 | 62 | 9 | 00:28<sup>+</sup> |
| **10** | 34 | M | BTLE | *RNS* | IIA | D | 14 | 40 | 4 | 15:10<sup>+</sup> |
| **11** | 69 | F | RTLE | RTL<sub>o</sub> | IIIA | D | 2 | 39 | 9 | 22:45<sup>#</sup> |
| **12\*** | 18 | M | LTLE, A<sup>1</sup> | LTL<sub>o</sub> | IIA | M | 11 | 161 | 7 | 14:43 |
| **13** | 22 | F | LTLE | *RNS* | IIIA | D | 2 | 75 | 5 | 05:03 |
| **14** | 53 | F | LTLE | LTL<sub>o</sub> | IA | D | 1 | 69 | 4 | 20:44<sup>+</sup> |
| **15** | 50 | F | BTLE | *RNS* | IIIA | D | 11 | 49 | 10 | 22:38<sup>+</sup> |
| **16\*** | 27 | M | BTLE, RTLS | *RNS* | IVB | D | 11 | 109 | 10 | 20:38<sup>+</sup> |

**Table 2:** Evaluation of ROCs of SOZ prediction by each estimated parameter. The following prediction metrics are shown across the *N* = 16 patients: Means and standard deviations of the Area Under the Curves (AUC), number of patients with AUC *>* 0.60 (Pred.), number of patients with AUC larger than the cutoff generated from randomly shuffled labels (Strong Pred.), number of patients with false positive rates less than 0.60 for true positive rates equal to 1 (FPR*<* 0.60), and number of patients with false positive rates less than 0.20 for true positive rates equal to 1 (FPR*<* 0.20). CC denotes prediction metrics by Clumping Coefficients. CV denotes prediction metrics by Coefficients of Variation. HR denotes prediction metrics by HFO Rate. 1 denotes prediction metrics estimated from parameters of Model 1. 2A denotes prediction metrics estimated from parameters of State A of Model 2. 2B denotes prediction metrics estimated from parameters of State B of Model 2.

| Parameter | AUC | Pred. | Strong Pred. | FPR < 0.60 | FPR < 0.20 |
| --- | --- | --- | --- | --- | --- |
| **CC1** | $0.81 \pm 0.18$ | 14 | 12 | 12 | 6 |
| **CC2A** | $0.82 \pm 0.14$ | 14 | 13 | 14 | 6 |
| **CC2B** | $0.75 \pm 0.21$ | 12 | 9 | 10 | 4 |
| **CV1** | $0.79 \pm 0.19$ | 12 | 10 | 12 | 8 |
| **CV2A** | $0.77 \pm 0.24$ | 11 | 10 | 13 | 7 |
| **CV2B** | $0.73 \pm 0.20$ | 11 | 8 | 12 | 4 |
| **HR1** | $0.67 \pm 0.26$ | 9 | 9 | 11 | 5 |
| **HR2A** | $0.70 \pm 0.30$ | 10 | 9 | 11 | 6 |
| **HR2B** | $0.63 \pm 0.25$ | 8 | 7 | 10 | 3 |

Long-term iEEG was recorded for all patients with high sample rates (minimum 2000 Hz, maximum 5000 Hz) in order to capture HFOs in the ripple band with high accuracy. Note that standard clinical sampling rates of 500 Hz and below may not be sufficient to capture ripples due to aliasing of digital signals. It is recommended that a sample rate of at least 250 * 2.5 = 625 Hz be used to capture ripple-band HFOs; the 2.5 multiplier is Engineer’s Nyquist given by Bendat and Piersol (2011). SOZ channels were identified by board-certified epileptologists as those with time courses indicative of seizure onset before propagation to other channels during any seizure captured via iEEG.

Channels were localized via coregistration of pre- and post-implantation magnetic resonance imaging (MRI) and/or post-implantation computed tomography (CT) as described by Stolk et al. (2017); Zheng et al. (2017); Helfrich et al. (2018); Stevenson et al. (2018). Each intracranial channel was classified as out-of-brain, within white matter, or within grey matter. If the location was on the boundary of the grey and white matter, it was labeled as white matter. If the location was near the edge of the brain, it was labeled as being outside the brain. We did not disaggregate by grey matter regions (hippocampus, amygdala, insular regions, neocortical regions, etc.), although other studies have described differing HFO rates between these regions (Blanco et al., 2011; Wang et al., 2017; Frauscher et al., 2018). Whenever possible, a channel within each strip or grid that was located within white matter was used as a reference. If no such information existed or was unclear from the localization, the channels were referenced to the average of all the channels on the grid or depth strip. The importance of coregistration was assessed with 6 of the 16 patients in which localization information was unavailable, and so we used data from all available iEEG electrodes. That is, we tested the robustness of our procedure to the absence of localization information that could have been used to exclude electrodes not placed in neural tissue or placed in white matter.

In two patients, scalp EEG, heart rate (electrocardiography; EKG), and eye movements (electrooculography, EOG) were concurrently recorded in order to extract sleep stage information over time in offline analysis. The data were then sleep staged using the software from Greer and Saletin (2011). Thirty-second epochs of data were classified as NREM slow wave sleep, REM, wakefulness, or artifact. This sleep staging was then compared to HFO model-found brain states, discussed later.

### 3.3 Automatic detection of high frequency oscillations during long-term recordings

Automatic detection of HFOs is now widely used, and the results of automatic detectors are comparable to that of visual detection (e.g. see Jacobs et al., 2018; Remakanthakurup Sindhu et al., 2020). We detected HFOs automatically in each channel of iEEG over the duration of each patient’s recording using the HFO detection software developed by Charupanit and Lopour (2017). This algorithm finds oscillations that are significantly larger than the amplitude noise floor in the 80 to 250 Hz frequency band. By iteratively generating a Poisson distribution of oscillation (“peak”) amplitudes, the detector can identify events with at least 4 consecutive high amplitude oscillations that exist in the tail of the rectified amplitude distribution (i.e. 8 rectified peaks). Specifically, we defined the threshold as peak amplitudes above the 95.8% percentile (i.e. *α* = 0.042, which was recommended by Charupanit and Lopour (2017)). Note that the estimation of the noise floor is adaptive and will change for each channel. We also allowed the noise floor to change every 5 minutes *within* each channel to account for non-stationarities, such as changes in state of vigilance and sleep stage.

To ensure that HFO rates were not affected by the occurrence of seizures, we analyzed only interictal HFOs that occurred at least 1 hour away from clinically-identified seizures. The resulting dataset had at least 4 hours of iEEG per patient, with a maximum of 25 hours for one patient, and a mean and standard deviation of 10 ± 5 hours across *N* = 16 patients (see **Table 1**). The original iEEG records contained overnight data. However our stipulations that the HFO counts used in analysis should both be consecutive and be at least 1 hour away from clinically-identified seizures resulted in diverse start times and coverage of these records. An example of the changing rate of HFOs from two channels within one patient is given in **Figure 1**.

**Figure 1:**
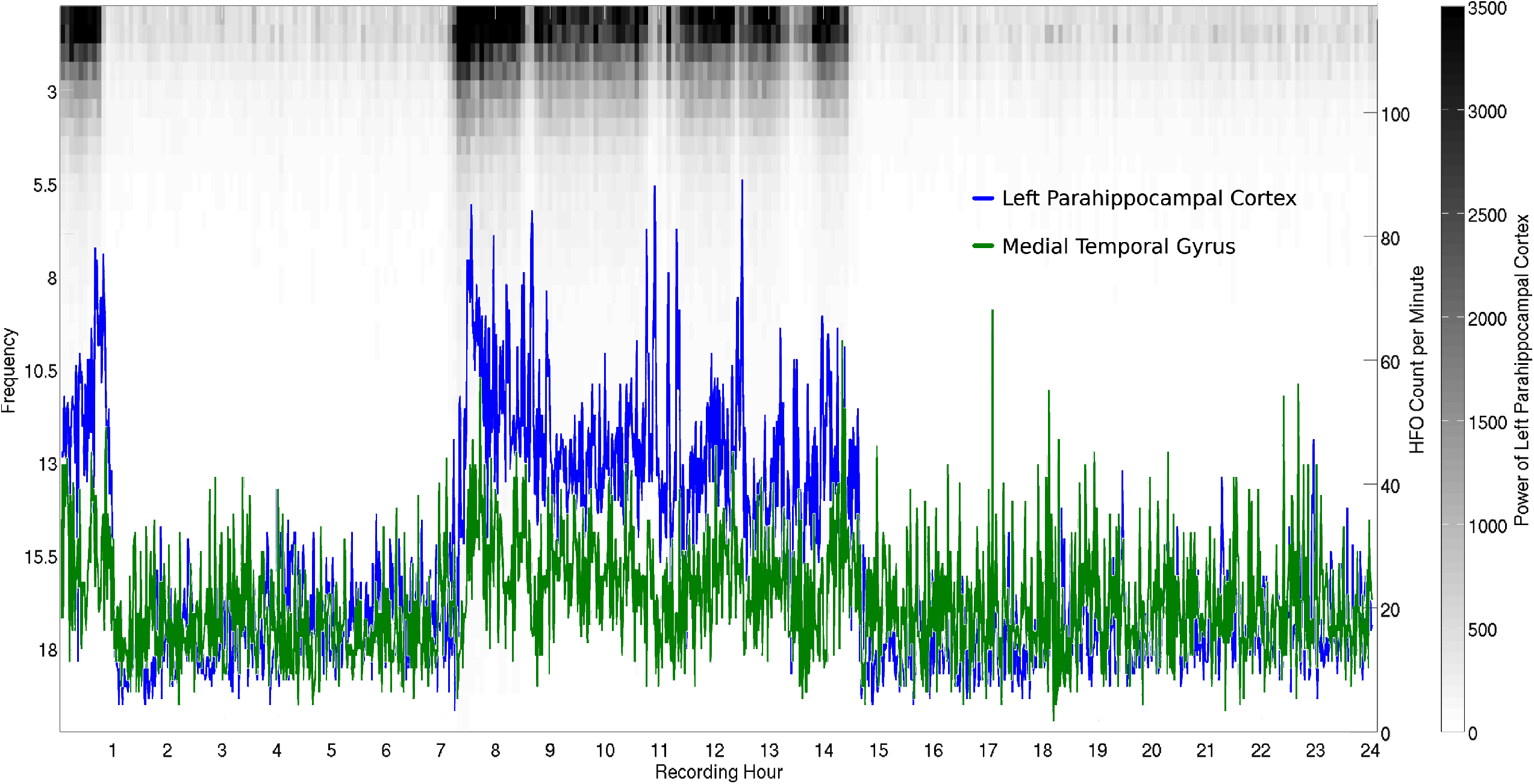
Modulation of HFO rate over 24 hours in one example patient. HFOs were automatically detected using the algorithm by Charupanit and Lopour (2017). Automated HFO counts per minute are shown from an iEEG channel in the left parahippocampal cortex (blue line) and an iEEG channel in the medial temporal gyrus (green line), with HFO counts per minute denoted by the right y-axis. These counts are overlayed on a grey-scale color map of low frequency band power (1-20 Hz) from the same left parahippocampal iEEG channel during the same 24 hour time period. Darker shading indicates higher power in the frequency band denoted by the left y-axis. HFO counts (blue and green lines) are modulated by the sleep-wake cycle (von Ellenrieder et al., 2017), as evidenced by their correlation with delta (1-4 Hz) frequency power (grey-scale color map). HFOs are typically analyzed during slow-wave sleep. However, with the method presented here, the data does not need to be visually sleep staged prior to classification of SOZ and non-SOZ channels.

### 3.4 Removing artifactual HFOs

We extended the Charupanit and Lopour (2017) algorithm by subsequently identifying and then removing detected HFOs that were likely artifact. This extension closely followed the “qHFO” algorithm of Gliske et al. (2016). That is, we sought to remove detected HFOs that (1) occurred in all channels simultaneously since the sources of these HFOs were likely spatially-broad electrical artifacts rather than localized neural generators, and (2) were falsely identified due to DC-shift artifacts that appeared as HFOs after bandpass filtering. Thus we first extracted HFOs using the algorithm by Charupanit and Lopour (2017), as discussed previously. We then calculated the common average across channels and reran the Charupanit and Lopour (2017) algorithm on this common average to identify cases of likely electrical artifacts. Then we found DC shifts in each channel by band passing the data from 850 to 990 Hz, calculating the line length of each 100 ms segment of data, and marking segments as DC shifts if they exceeded a threshold of 4 standard deviations above the mean line length over the previous 5 seconds (again calculated in 100 ms segments). HFO occurrences for each iEEG channel that overlapped in time with either artifactual HFOs found in the common average or DC shifts detected in that channel were removed from further analysis.

### 3.5 Assuming Negative Binomial processes

It has previously been shown that pathological, interictal HFOs can be modulated by high amplitude, low frequency background activity during sleep (Kerber et al., 2014; Frauscher et al., 2015; von Ellenrieder et al., 2016; Motoi et al., 2018). Therefore the typical model for count data, a Poisson process in which the variance must equal the mean rate over time (Cook, 2009), may not accurately describe HFO count data in general. While we did not directly measure phase-amplitude coupling of HFOs to slow rhythms, we estimated the variance of the HFOs over time within fitted models. “Overdispersion” occurs when there is greater variability in the data (e.g. HFO count data) than is expected by a Poisson process. Overdispersion can be due to the “clumping” of HFOs in close temporal proximity to one another, such that there are bursts of HFOs occurring in time followed by relatively quiet periods without many HFOs (see bottom plot of **Figure 2**). Thus we might expect overdispersion to be predictive of SOZ based on the previous literature. We estimated parameters in hierarchical models that provide inference as to whether HFOs occur in patterns in which they are “clumped” together.

**Figure 2:**
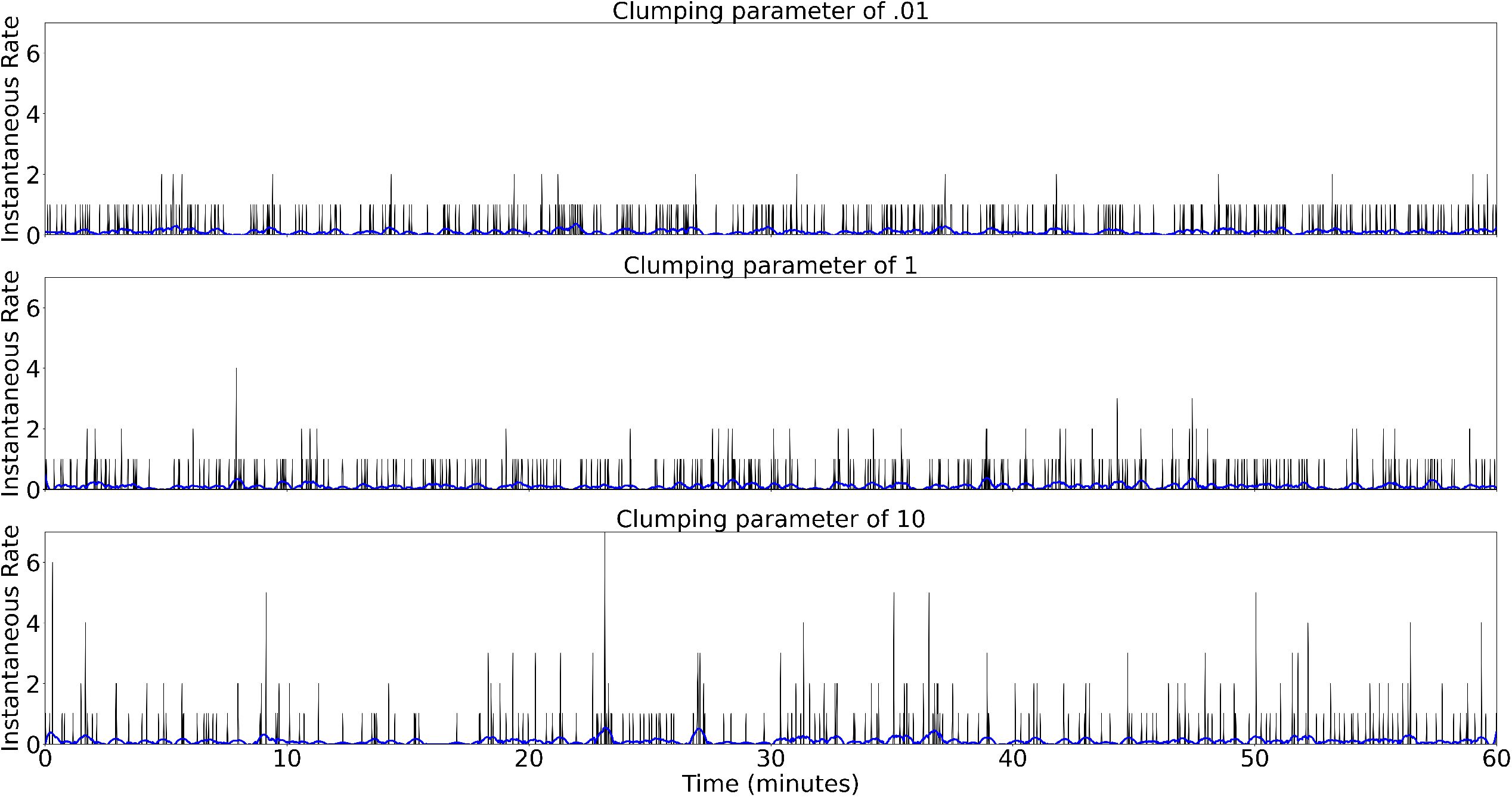
Top: Simulated instantaneous HFO rates (per second) from a Negative Binomial process with a small clumping coefficient (*ζ* = 0.01, near a Poisson process) over a 60 minute period. Electrodes that contained Negative Binomial processes with small clumping coefficients were found to be predictive of SOZ. Middle: Simulated instantaneous HFO rates from a Negative Binomial Process with some clumping (*ζ* = 1). Bottom: Simulated instantaneous HFO rates from a Negative Binomial Process with significant clumping (*ζ* = 10). All simulations had a HFO rate of *λ* = .1 per second.

A Negative Binomial process is a description of count data that can account for overdispersion (Cook, 2009) and can be viewed as a relaxation of a restriction that HFO counts must follow a strictly Poisson process. In order to characterize the temporal dynamics of HFOs, we fit the data to Negative Binomial models of count data per second. That is, we automatically counted the number of HFOs from our qHFO algorithm for each iEEG electrode and each patient per second. These counts per second were then assumed to be generated from a Negative Binomial process whose parameters could change every 5 minutes. We chose a 5-minute window in order to measure changes with high time resolution over multiple hours while also keeping enough observations to accurately estimate parameters of the Negative Binomial distribution of count data. This resulted in 60 * 5 = 300 HFO count observations to estimate parameters of the Negative Binomial process per 5 minutes.

The Negative Binomial distribution has two common parameterizations. In the parameterization we used in this study, the Negative Binomial distribution gives a number of “failures” (e.g. number of HFOs within a given second) before *η* “successes” where *θ* is the probability of a success (e.g. the probability that no HFOs occur) (Plummer, 2003; Cook, 2009). Note that *η* is not restricted to integers. To aid SOZ prediction, we transformed the two parameters of the Negative Binomial distribution *θ* and *η* to create three parameters: (1) the rate of HFOs per second *λ*, (2) a “clumping coefficient” (CC, *ζ*), defined as the inverse of *η*, and (3) the coefficient of variation (CV, *γ*), defined as the ratio of the standard deviation of counts over the rate of HFOs. Note that as *η* goes to positive infinity, the clumping coefficient *ζ* goes to zero and the Negative Binomial distribution approaches a Poisson distribution (Cook, 2009). The top plot of **Figure 2** shows an example of a small clumping coefficient. Large clumping coefficients *ζ* indicate that HFOs are more likely to occur immediately following other HFOs (see bottom plot of **Figure 2**). CV much greater than 1 (*γ >>* 1) also indicate temporal clumping of HFOs, i.e. a CV greater than one indicates that the variance is greater than the mean rate. CV much less than 1 (*γ <<* 1) indicate oscillatory dynamics, i.e. a CV less than one indicates a low variance relative to the mean rate. A CV near 1 (in addition to a clumping coefficient near zero, *ζ* ≈ 0) indicates that the process is more Poisson-like. The HFO rate, CC, and CV were included in the model to gauge the predictive ability of each parameter to inform the location of the SOZ.

To illustrate the concept of clumping, we simulated Negative Binomial processes. Specifically, we simulated three different clumping coefficients, *ζ* = 0.01, *ζ* = 1, and *ζ* = 10, with an HFO rate parameter of *λ* = .1 per second over a 60 minute period. Note that as the clumping coefficient approaches zero, *ζ* → 0, the number of “successes” approaches positive infinity, *η* → ∞, and the Negative Binomial process with rate *θ* approaches a Poisson process with rate *θ*. Thus *ζ* = 0.01 approximates a Poisson process. In the simulation, we disregarded the shape and duration of the HFOs themselves and instead simulated the number of HFOs per second per electrode. The simulation results are shown in **Figure 2**.

### 3.6 Hierarchical Negative Binomial models

To automatically obtain HFO dynamics across channels within each patient, we fit models to the HFO count data using Bayesian methods with JAGS (Just Another Gibbs Sampler). JAGS can easily sample from complex models using Markov Chain Monte Carlo (MCMC) (Plummer, 2003) via the pyjags Python package (Miasko, 2017). Specifically, we fit models of HFO counts per second *t* which contained parameters of HFO dynamics and hierarchical parameters. We then derived estimates of HFO rates *λ*, clumping coefficients *ζ*, and coefficients of variation *γ*. Hierarchical parameters were included in the model to encourage stable parameter estimates across the 5-minute windows *w*, as we expected iEEG channels to have somewhat consistent temporal dynamics. Hierarchical distributions of HFO parameters allow parameters to “shrink” towards the mean HFO parameters, which improves estimation of these parameters in the presence of outliers (Gelman et al., 2013; Boehm et al., 2018). We experimented with different hierarchical models with Poisson and Negative Binomial baselikelihoods describing HFO counts per second *t* without assessing SOZ prediction. Many models’ parameters would not converge to stable posterior distributions, either due to an excess of hierarchical parameters which caused the models to be unidentifiable or due to a complexity in the parameter space that the splice sampling in JAGS has difficulty sampling. We present parameter fitting results and SOZ prediction from two hierarchical Negative Binomial models in this paper, first without (Model 1) and then with (Model 2) an undetermined mixture of distributions over time. We also present results from a mixture model of Poisson distributions (Model 3) in the Supplementary Materials.

In Model 1, we assumed that the HFO dynamics were described by a Negative Binomial distribution and that these dynamics could change per 5-minute window. We also included hierarchical distributions such that the HFO rate *λ*, measured in 5-minute windows *w*, was described by a normal distribution with a mean HFO rate parameter *μ*_(*λ*)_ with some standard deviation *σ*_(*λ*)_ for each iEEG channel *e*. Similarly, the clumping coefficient *ζ* was described by a normal distribution with mean clumping coefficient *μ*_(*ζ*)_ with some standard deviation *σ*_(*ζ*)_ across the 5-minute windows for each iEEG channel *e*. The coefficient of variation *γ* was derived from the ratio of the standard deviation to the mean of the Negative Binomial distribution, and it was not described by a hierarchical distribution of parameters.

Model 1 was fit to the HFO count data of each patient separately and is given by the following likelihood distribution, parameter relationship equations, hierarchical distributions, and prior distributions:

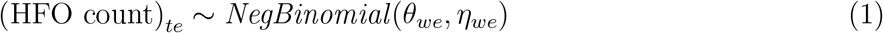

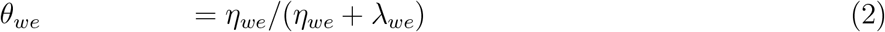

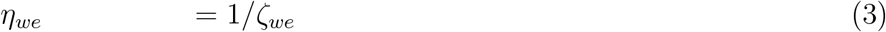

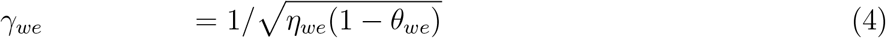

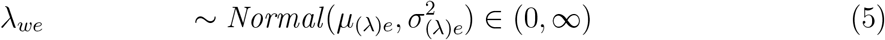

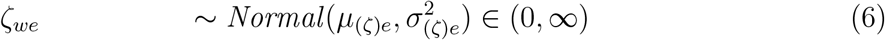

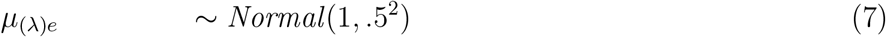

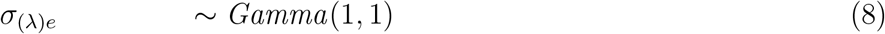

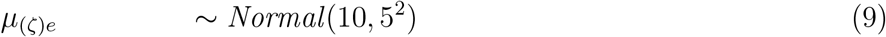

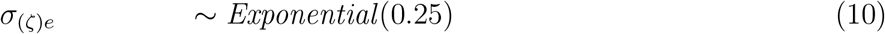

### 3.7 Assuming mixtures of Negative Binomial distributions

Based on previous research (Dümpelmann et al., 2015; von Ellenrieder et al., 2016, 2017), we assumed that HFO rates would be a function of the state of vigilance and sleep stage. This could be seen when the HFO rates were plotted over time, as increased rates correlated with increased delta (1-4 Hz) power, which is generally indicative of slow-wave sleep (see **Figure 1**). We also suspected that HFO rates might change based on the cognitive brain state of the patient. For these two reasons we allowed another hierarchical model, Model 2, to automatically identify the states inherent in the data.

Mathematically, in Model 2 we assumed that HFO counts per second for each channel were distributed from a mixture model of Negative Binomials. That is, we assumed that each channel contained multiple distributions of HFO counts (one distribution per state *k*), with the representative state changing over time. We enforced the restriction that a change in state caused the distributions from all channels to change at the same time. Thus, we assumed that the brain’s state of vigilance or sleep changed each channel’s HFO dynamics simultaneously, although the parameter values for each channel could change in different ways. We initially constrained the number of possible brain states *k* to 2-4 per channel, switching at most every five minute window *w* during the recordings. After initial model fitting experiments, discussed above, we constrained the number of Negative Binomial mixtures to be *two* per channel.

Model 2 was given by the following equations:

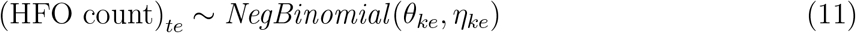

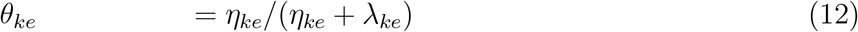

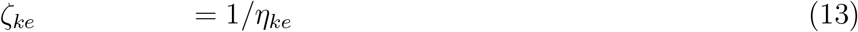

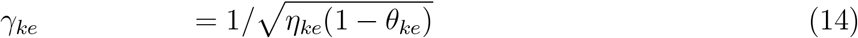

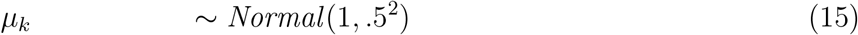

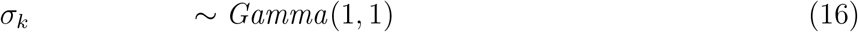

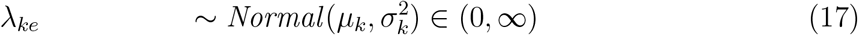

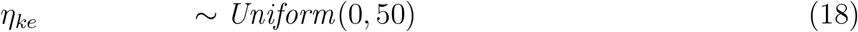

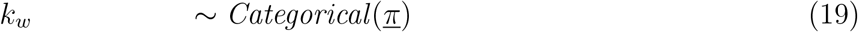

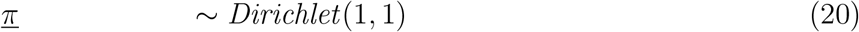

### 3.8 Solving model convergence issues

Each model was fit using Markov Chain Monte Carlo in JAGS with six chains of 5,200 samples each. This was performed in parallel with 200 burn-in samples and a thinning parameter of 10. This procedure resulted in (5,200 − 200)*/*10 = 500 posterior samples from each chain for each parameter. We kept all posterior samples from each chain to assess posterior distributions from Model 1. We confirmed that this model fitting procedure produced useful parameter estimates in simulation (see Supplementary Methods on simulated Negative Binomial processes).

However, the Markov chains resulting from Model 2 suggested that this model may not easily converge to posterior distributions, depending upon the initial conditions and given HFO count data. Obtaining model convergence is often difficult with mixture-modeling in general, and it was not easily solved when assuming a certain number of brain states in our modeling work presented here. For those patients whose data did not converge when assuming two brain states, two-state models were enforced by removing non-converging Markov Chains in order to achieve convergence across all kept chains. Out of the 6 Markov Chains for each model, a chain was removed if (1) its time course over samples did not converge to a one-peaked posterior distribution, and (2) if the chain did not converge to the remaining majority of other chains (if applicable). We also calculated the Gelman-Rubin statistic, 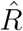, for each parameter; this compares the estimated between-chain and within-chain variances (Gelman and Rubin, 1992). Six patients’ data had Model 2 converge in all six chains, five patients’ data had Model 2 converge in five of six chains, one patient’s data had Model 2 converge in half the chains, three patient’s data had Model 2 converge in two chains, and one patient’s data had Model 2 “converge” with one chain. The posterior samples from each chain of 500 samples were combined to form one posterior sample between 500 and 3,000 samples for each parameter in each model. Note that removing chains is an unorthodox method in Bayesian analysis, and *does not strictly guarantee model convergence*. However, this procedure enabled better prediction of the SOZ than Model 1 in some patients, especially when using the clumping coefficient and coefficient of variation as shown in **Figures 4** and **5**. However the non-convergence results may imply that a two-state model is *not* sufficient to *describe all HFO dynamics*.

### 3.9 Classification of SOZ and non-SOZ channels

We used estimates of the HFO rate parameter, the clumping coefficient (CC), and the coefficient of variation (CV) obtained from the posterior distributions of Model 1 to classify channels as SOZ or non-SOZ. Specifically, we took the average across time windows *w* of the posterior medians of *λ, ζ*, and *γ* to generate estimates for each iEEG channel *e* with each patient’s data. Note that we used the mean of median posteriors from Model 1 as the estimates for rate and the CC instead of the hierarchical mean parameters of HFO rate *μ*_(*λ*)*e*_ and CC *μ*_(*ζ*)*e*_, and we confirmed that the mean of median posteriors were reflective of true mean HFO rates and CCs in simulation (see **Figure 8**). In contrast, all HFO parameters from Model 2 were estimated by the medians of posterior distributions for each brain state *k*. After finding parameter estimates the brain states were sorted by the average amount of standardized mean delta (1-4 Hz) power across all iEEG channels and 5-minute windows used in the model, and we will refer to them as brain state A (brain state with higher mean delta power) and brain state B (brain state with lower mean delta power), see **Figure 7**.

We built Receiver Operating Characteristic (ROC) curves by varying the cutoff values for classification. ROC curves show the trade-off between true positives (channels identified as SOZ by clinicians that are also labeled as candidate SOZ channels by the cutoff value) versus false positives. Clinicians and researchers may find ROC curves useful because of the possible trade-offs between resecting or not-resecting some identified SOZ tissue. These ROC curves show the overall accuracy of our prediction over a continuum of possible cutoff values. One ROC curve was created for each patient and each HFO parameter. The Area Under the Curve (AUC) was also calculated for each patient and parameter by integrating the ROC curves. AUCs were viewed as an overall measure of predictability of each parameter for each patient. We evaluated AUC values for each parameter within in each patient by (1) deeming AUC *>* 0.60 as “predictive” and (2) comparing performance of ROC curves based on real SOZ and non-SOZ labels to ROC curves based on randomly shuffled labels. That is, we randomly shuffled SOZ and non-SOZ labels of each channel used in the modeling without replacement 1000 times and calculated 1000 fake AUCs values. We then ordered the fake AUCs from smallest to largest for each parameter and patient and found the 950th value (95% of the reshuffled samples) to use as an AUC cutoff. If the AUC was larger than this AUC cutoff, the AUC was deemed “strongly predictive”. We also built Precision-Recall curves (Davis and Goadrich, 2006) that are shown and explained in the Supplementary Materials. However, Precision-Recall curves cannot easily be compared across patients due to different baseline ratios of SOZ to non-SOZ channels (see **Table 1**).

## Data and code availability

Automatically identified HFO counts, standardized delta (1-4 Hz) power, channel localizations, and samples from posterior distributions for Models 1-3 are available upon request and at https://doi.org/10.6084/m9.figshare.12385613. MATLAB, Python, and JAGS data extraction and analysis code are available at https://osf.io/3ephr/ and in the following repository https://github.com/mdnunez/sozhfo (as of June 2020 with a major update in August 2021).

## 5 Results

### 5.1 Small clumping coefficients are predictive of SOZ

The data and modeling show that small clumping coefficients (CC) are predictive of SOZ, as judged by evaluating parameter estimates of CC from both models. We will refer to clumping coefficients estimated by Model 1 as “CC1”, the clumping coefficients estimated by Model 2 in brain state A as “CC2A”, and the clumping coefficients estimated by Model 2 in brain state B as “CC2B”. The mean and standard deviation of the CC1 AUCs across patients were 0.81 ± 0.18, while the same statistics derived from CC2A and CC2B were 0.82 ± 0.14 and 0.75 ± 0.21 respectively. The data of 14 of 16 patients yielded CC1 and CC2A that we deemed predictive of SOZ (AUC *>* 0.60), while the data of 12 patients yielded CC2B that were deemed predictive. The ROC curves and distribution of AUCs for the CC parameter are shown in **Figure 3**. All summary ROC evaluation statistics are given in **Table 3**.

**Table 3:**
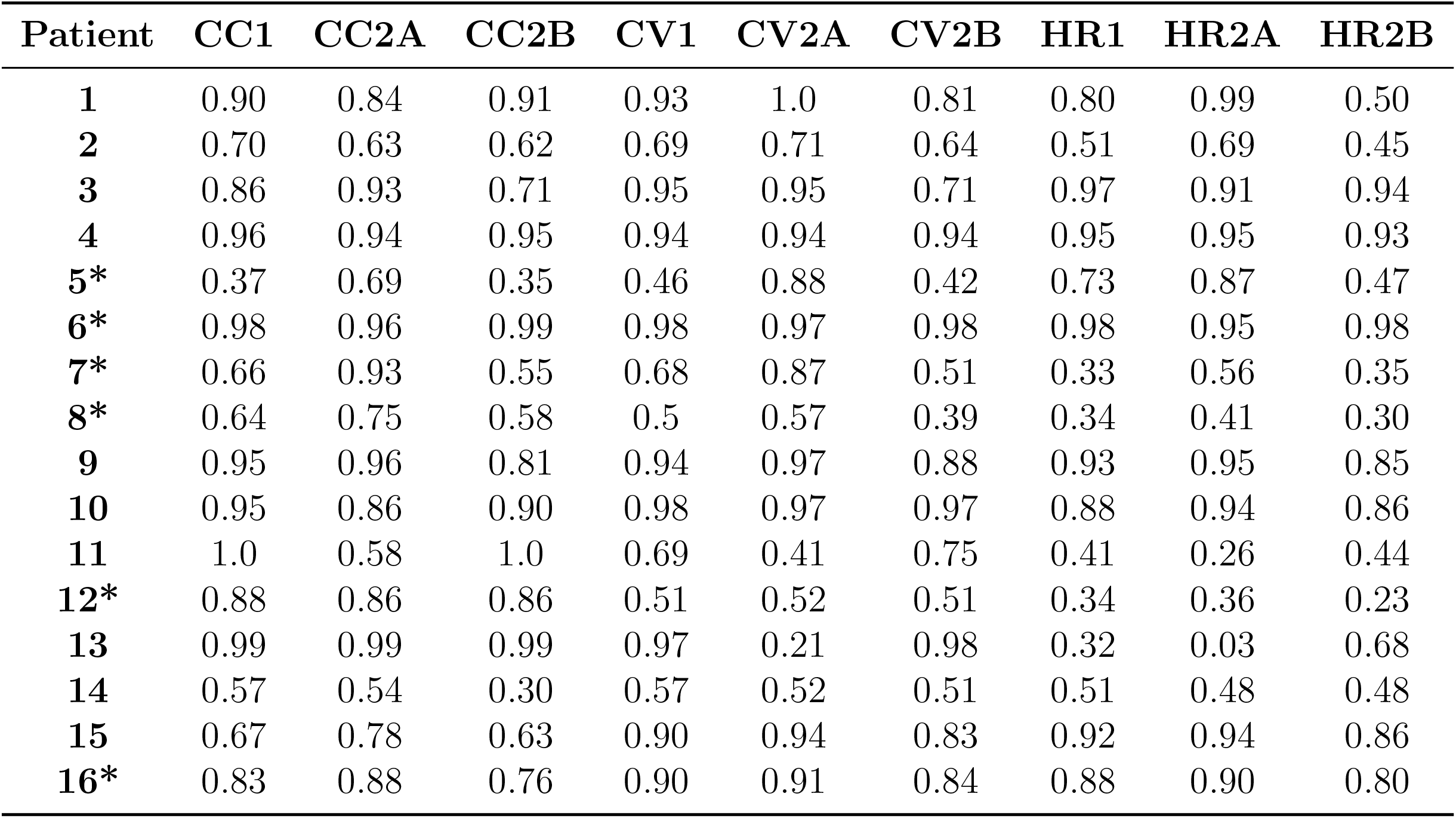
Evaluation of ROCs by Area Under the Curves (AUC) for each patient. CC denotes prediction metrics by Clumping Coefficients. CV denotes prediction metrics by Coefficients of Variation. HR denotes prediction metrics by HFO Rate. 1 denotes prediction metrics estimated from parameters of Model 1. 2A denotes prediction metrics estimated from parameters of State A of Model 2. 2B denotes prediction metrics estimated from parameters of State B of Model 2. * denotes patients for whom all channels were used to fit data to the mixture models, while we included only grey matter localized channels for the other patients.

| Patient | CC1 | CC2A | CC2B | CV1 | CV2A | CV2B | HR1 | HR2A | HR2B |
| --- | --- | --- | --- | --- | --- | --- | --- | --- | --- |
| **1** | 0.90 | 0.84 | 0.91 | 0.93 | 1.0 | 0.81 | 0.80 | 0.99 | 0.50 |
| **2** | 0.70 | 0.63 | 0.62 | 0.69 | 0.71 | 0.64 | 0.51 | 0.69 | 0.45 |
| **3** | 0.86 | 0.93 | 0.71 | 0.95 | 0.95 | 0.71 | 0.97 | 0.91 | 0.94 |
| **4** | 0.96 | 0.94 | 0.95 | 0.94 | 0.94 | 0.94 | 0.95 | 0.95 | 0.93 |
| **5\*** | 0.37 | 0.69 | 0.35 | 0.46 | 0.88 | 0.42 | 0.73 | 0.87 | 0.47 |
| **6\*** | 0.98 | 0.96 | 0.99 | 0.98 | 0.97 | 0.98 | 0.98 | 0.95 | 0.98 |
| **7\*** | 0.66 | 0.93 | 0.55 | 0.68 | 0.87 | 0.51 | 0.33 | 0.56 | 0.35 |
| **8\*** | 0.64 | 0.75 | 0.58 | 0.5 | 0.57 | 0.39 | 0.34 | 0.41 | 0.30 |
| **9** | 0.95 | 0.96 | 0.81 | 0.94 | 0.97 | 0.88 | 0.93 | 0.95 | 0.85 |
| **10** | 0.95 | 0.86 | 0.90 | 0.98 | 0.97 | 0.97 | 0.88 | 0.94 | 0.86 |
| **11** | 1.0 | 0.58 | 1.0 | 0.69 | 0.41 | 0.75 | 0.41 | 0.26 | 0.44 |
| **12\*** | 0.88 | 0.86 | 0.86 | 0.51 | 0.52 | 0.51 | 0.34 | 0.36 | 0.23 |
| **13** | 0.99 | 0.99 | 0.99 | 0.97 | 0.21 | 0.98 | 0.32 | 0.03 | 0.68 |
| **14** | 0.57 | 0.54 | 0.30 | 0.57 | 0.52 | 0.51 | 0.51 | 0.48 | 0.48 |
| **15** | 0.67 | 0.78 | 0.63 | 0.90 | 0.94 | 0.83 | 0.92 | 0.94 | 0.86 |
| **16\*** | 0.83 | 0.88 | 0.76 | 0.90 | 0.91 | 0.84 | 0.88 | 0.90 | 0.80 |

**Figure 3:**
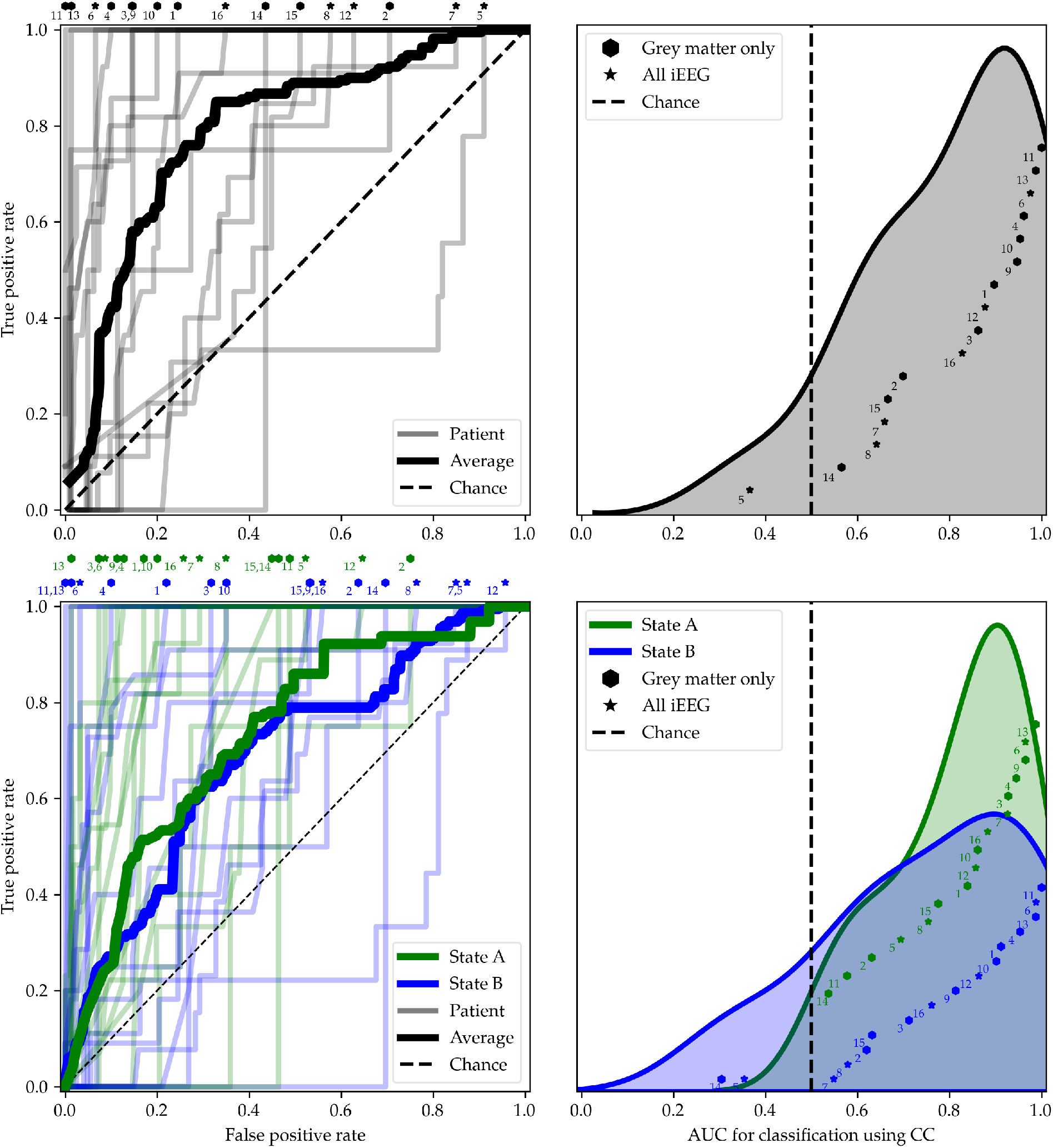
SOZ prediction based on the clumping coefficients (CC) estimated by Model 1 (Top) and Model 2 (Bottom). (Left) ROC curves when using **small** values of the CC to identify SOZ channels. ROC curves for individual patients (*N* = 16) are displayed using fine lines, and the average is shown in bold. The bold dashed line indicates an ROC at chance prediction. Data points on the top of these two plots indicate the false positive rate (FPR) for which the true positive rate (TPR) is 1, with patient labels to compare across plots. From Model 2, the brain state with the most delta (1-4 Hz) power in each patient was labeled State A (green lines), while the other model-found brain state was labeled State B (blue lines). (Right) Distribution of AUC values based on CC for each patient. The exact AUC values are denoted as hexagons or stars on the x-axis, while the shaded distributions are a density approximation from *N* = 16 values. Hexagons denote patients where the analysis was performed exclusively on grey matter channels, while stars denote patients for which all channels were included. Each point is labeled with the corresponding patient number, and the y-values are sorted by AUC; however, the y-values have no other meaning. The bold dashed lines indicate an AUC of 0.5.

Importantly, the prediction of SOZ by the clumping coefficient does not clearly depend upon localization of grey matter channels using CT and MRI scans (and exclusion of all other channels from the analysis). For instance, small CC2A were predictive of SOZ (AUC *>* 0.60) in all 6 patients for which we did not exclude channels that were outside the brain.

By combining all channels across all patients, we can also obtain information about the general predictability of SOZ using these parameters. The number of channels used in the models varied by patient (minimum of 41, maximum of 172, and a mean and standard deviation of 87 ± 39 channels across *N* = 16 patients, see **Table 1**), and the number of SOZ channels also varied by patient (minimum of 1, maximum of 14, and a mean and standard deviation of 7 ± 4). However, combining channels across patients (total channel count of 1391) provides estimates of the cutoff values for these parameters that could be used to identify SOZ channels during interictal periods. The aggregate ROC curves for the clumping coefficients estimated using Model 1 and Model 2 are shown in **Figure 4**. Across all *N* = 16 patients, when CC2A less than or equal to 1 (*ζ* ≤ 1) were treated as indicative of the SOZ, the FPR was only 0.31 across all channels, with a corresponding TPR of 0.86. Note that CC1≤ 1 estimates would result in a FPR of 0.22 and a TPR of 0.70. A TPR of 1 was achieved by treating all CC2A less than or equal to 2.34 (*ζ* ≤ 2.34) as indicative of the SOZ, although this resulted in a FPR of 0.62. The CC2B were not as informative of SOZ.

**Figure 4:**
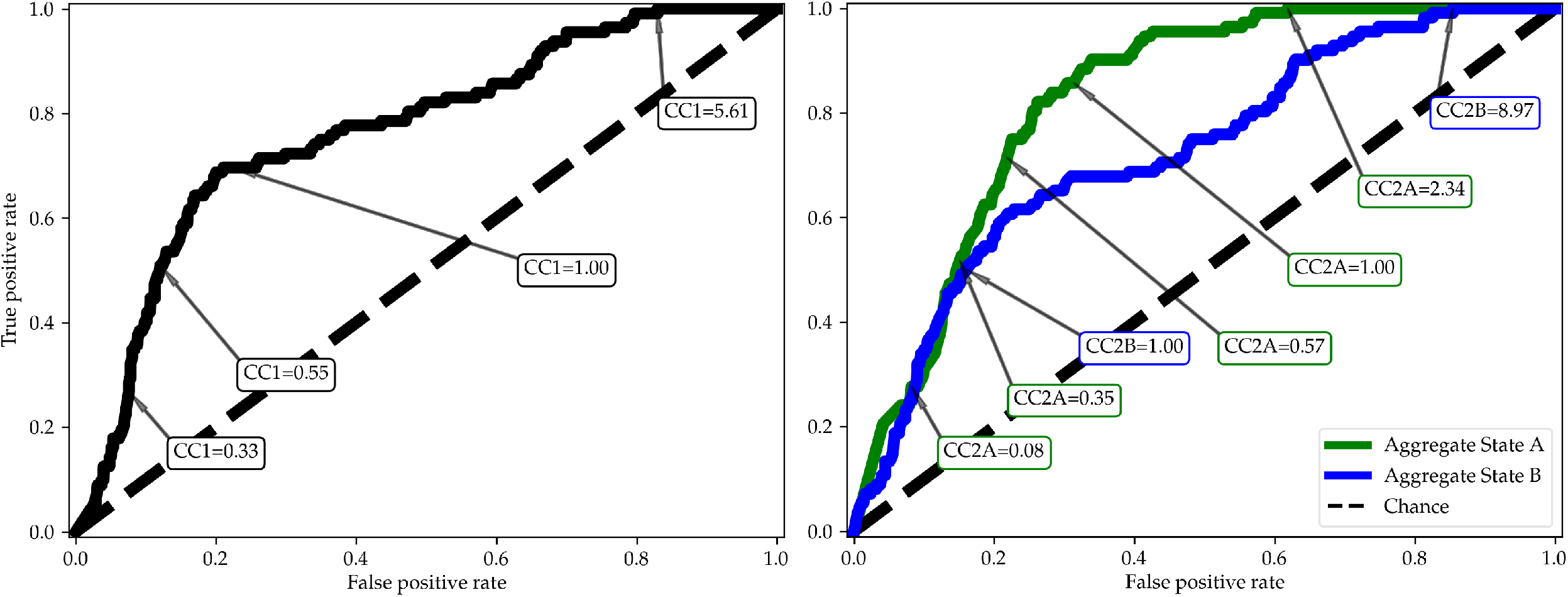
The aggregate ROC curves for all included channels (total channel count of 1391) across all patients (*N* = 16) when using **small** values of the HFO clumping coefficient (CC) estimates to identify SOZ channels. A few representative cutoff CC values are shown in the text boxes, such that values smaller than the CCs in text boxes generated the indicated points on the ROC curves across all included patients. (Left) Aggregate ROC curve generated from CC estimates using Model 1 (CC1). (Right) Aggregate ROC curve generated from CC estimates using Model 2. The brain state with the most delta (1-4 Hz) power in each patient was labeled brain state A while the other model-found brain state was labeled brain state B. The green curve was generated from all CCs for each channel from all patients’ model-found brain states A (CC2A). The blue curve was generated from all CCs for each channel from all patients’ model-found brain states B (CC2B). The bold dashed lines indicate an AUC of 0.5.

### 5.2 Coefficients of variation less consistently predict SOZ

The ability of the coefficient of variation (CV) to predict SOZ was similar, but slightly less consistent, than the clumping coefficients. We will refer to CV estimated by Model 1 as “CV1”, the CV estimated by Model 2 in brain state A as “CV2A”, and the CV estimated by Model 2 in brain state B as “CV2B”. The ROC curves and distribution of AUCs for the CV parameter are shown in **Figure 5**. The mean and standard deviation of the CV1 AUCs across patients were 0.79 ± 0.19, while the same statistics derived from CV2A and CV2B were 0.77 ± 0.24 and 0.73 ± 0.20 respectively. ROC evaluation statistics based on prediction by CV are shown in **Table 3**.

**Figure 5:**
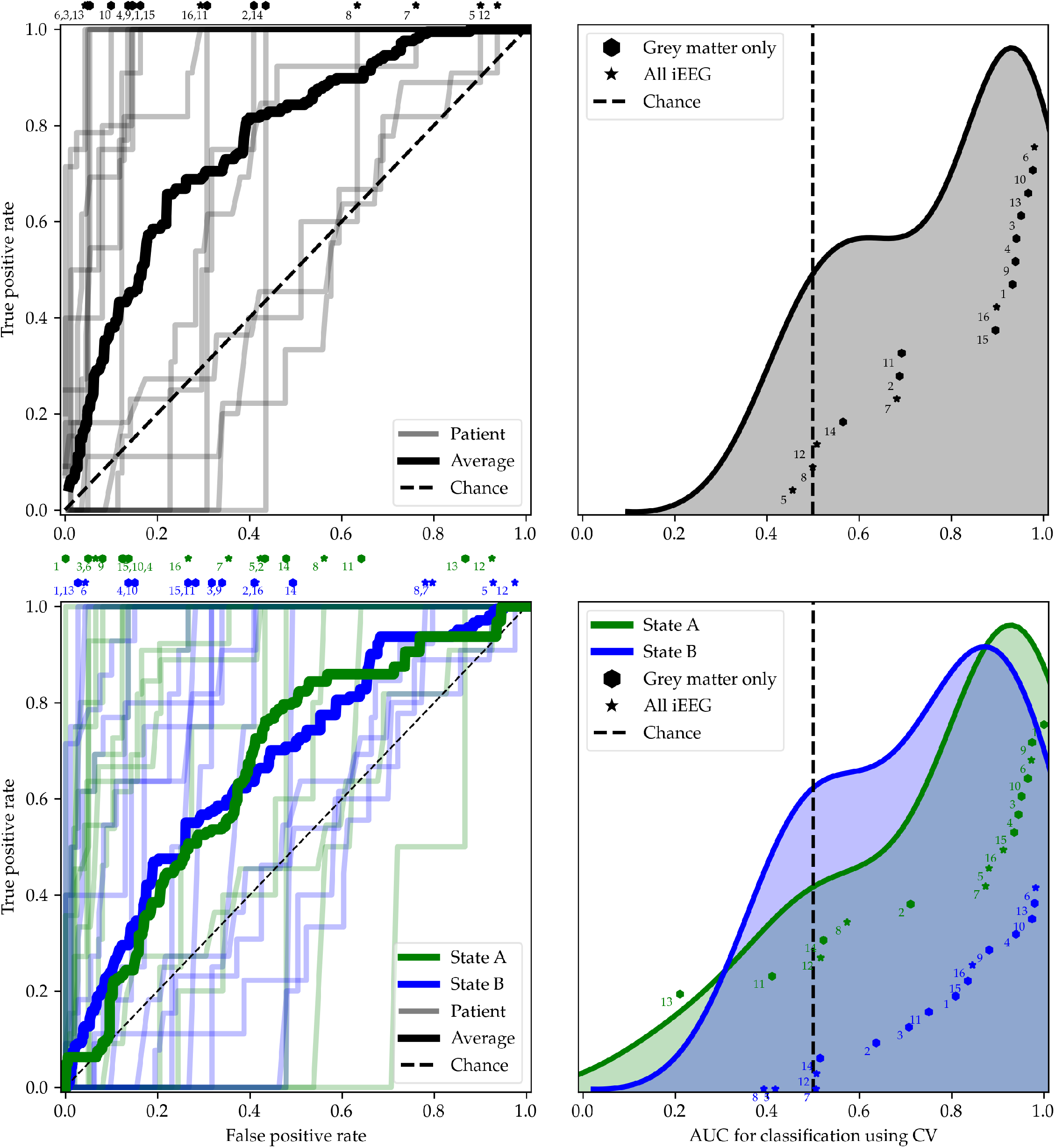
SOZ prediction based on the coefficients of variation (CV) estimated by Model 1 (Top) and Model 2 (Bottom). (Left) ROC curves when using **small** values of the CV to identify SOZ channels. (Right) Distribution of AUC values based on CV for each patient. Readers are referred to the caption of **Figure 3** for a detailed description of the plot elements.

### 5.3 Prediction of SOZ using HFO rate is not consistent across patients

In some patients, the HFO rate could be used to identify the SOZ channels in different states with success rates similar to the CC and CV parameters. However, large HFO rates were not predictive of SOZ in some patients. We will refer to HFO rates estimated by Model 1 as “HR1”, the HFO rates estimated by Model 2 in brain state A as “HR2A”, and the HFO rates estimated by Model 2 in brain state B as “HR2B”. The ROC curves and distribution of AUCs for the HFO rates parameter are shown in **Figure 6**. The mean and standard deviation of the HR1, HR2A, and HR2B AUCs across patients were 0.67 ± 0.26, 0.70 ± 0.30 and 0.63 ± 0.25 respectively. ROC evaluation statistics based on prediction by HFO rate are shown in **Table 3**.

**Figure 6:**
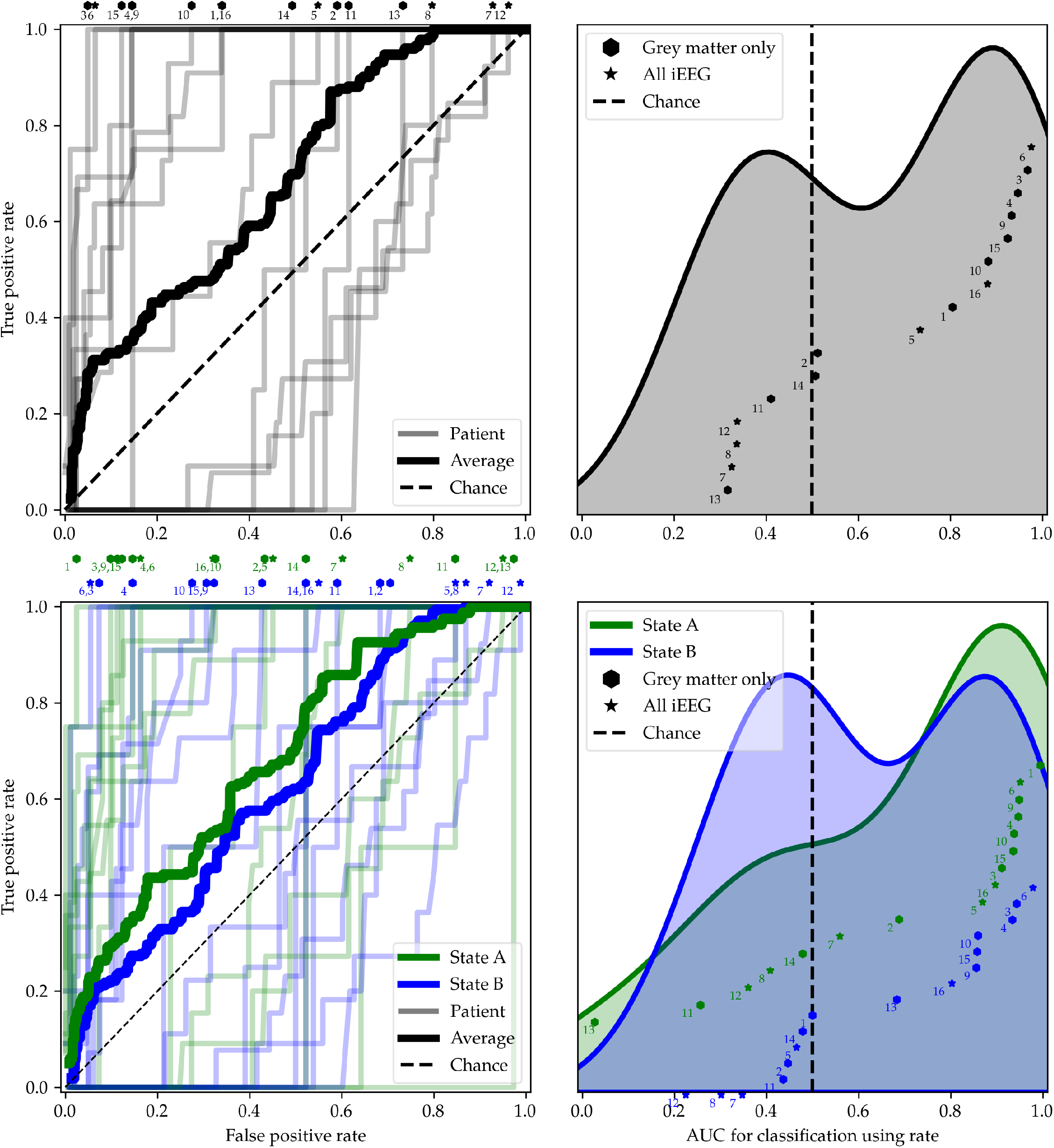
SOZ prediction based on the HFO rate estimated by Model 1 (Top) and Model 2 (Bottom). (Left) ROC curves when using **large** values of the HFO rate to identify SOZ channels. (Right) Distribution of AUC values based on HFO rate for each patient. Readers are referred to the caption of **Figure 3** for a detailed description of the plot elements.

### 5.4 Model-derived brain states correspond to sleep and wakefulness

The Negative Binomial mixture model (Model 2) automatically separated windows of time into two brain states based on the HFO dynamics in all channels. Brain state labels were thus influenced by the HFO dynamics from all channels simultaneously. We found that the brain state labels of converged Markov chains tracked known sleep/wake dynamics.

In two patients whose data were sleep staged manually using concurrent scalp EEG, the two HFO model-derived brain states blindly separated slow wave sleep (i.e. NREM sleep stages 1, 2 and 3) from all other states (REM and wakefulness) as shown in the lower two panels of **Figure 7**. In Patient 15, the congruence between the visually sleep staged data and the HFO model-derived brain states from Model 2 was 89.2%, with NREM sleep being correctly identified by the model 90.6% of the time and the other states being correctly identified 83.3% of the time. In Patient 16, the congruence between the sleep staged data and the model-derived brain states was 91.7%, with all states being correctly identified 91.7% of the time by the model.

We did not have concurrent EEG, EKG, and EOG in the other patients to evaluate the correspondence between sleep stages and model-derived brain states. However, we could evaluate how well standardized delta power (1-4 Hz), averaged across electrodes, corresponded to the two model-derived brain states, as a proxy for sleep staged data. In half of the patients (8/16), we found found that delta power was significantly different in the two states (*p <* .001) using both ANOVA and Kruskal-Wallis tests using cutoff *α* = .001 (see last column of **Table 1**). An additional patient had a significant Kruskal-Wallis test (*p* = .006) using cutoff *α* = .01, with a small ANOVA p-value (*p* = .011). And one more patient had a significant Kruskal-Wallis test (*p* = .025) using a cutoff of *α* = .05. Of the six patients without any indication of significant differences in delta power between the two model-derived brain states, four did not have localization information to enable exclusion of electrodes outside the brain prior to the standardized mean delta calculation. Only two of 10 patients for whom we included localization information did not show evidence for model-derived brain states consistent with changes in delta power. Note that we explored removing delta power outliers across channels (with various cutoffs of 1, 2, 3, and 4 standard deviations above the mean power across electrodes in each 5-minute time window) in a post-hoc analysis. While this did switch the brain state labels for four patients’ brain states (patients 3, 6, 7, and 11), the delta power in these patients still did not clearly differentiate between the two states (see the second panel of **Figure 7** for an example).

**Figure 7:**
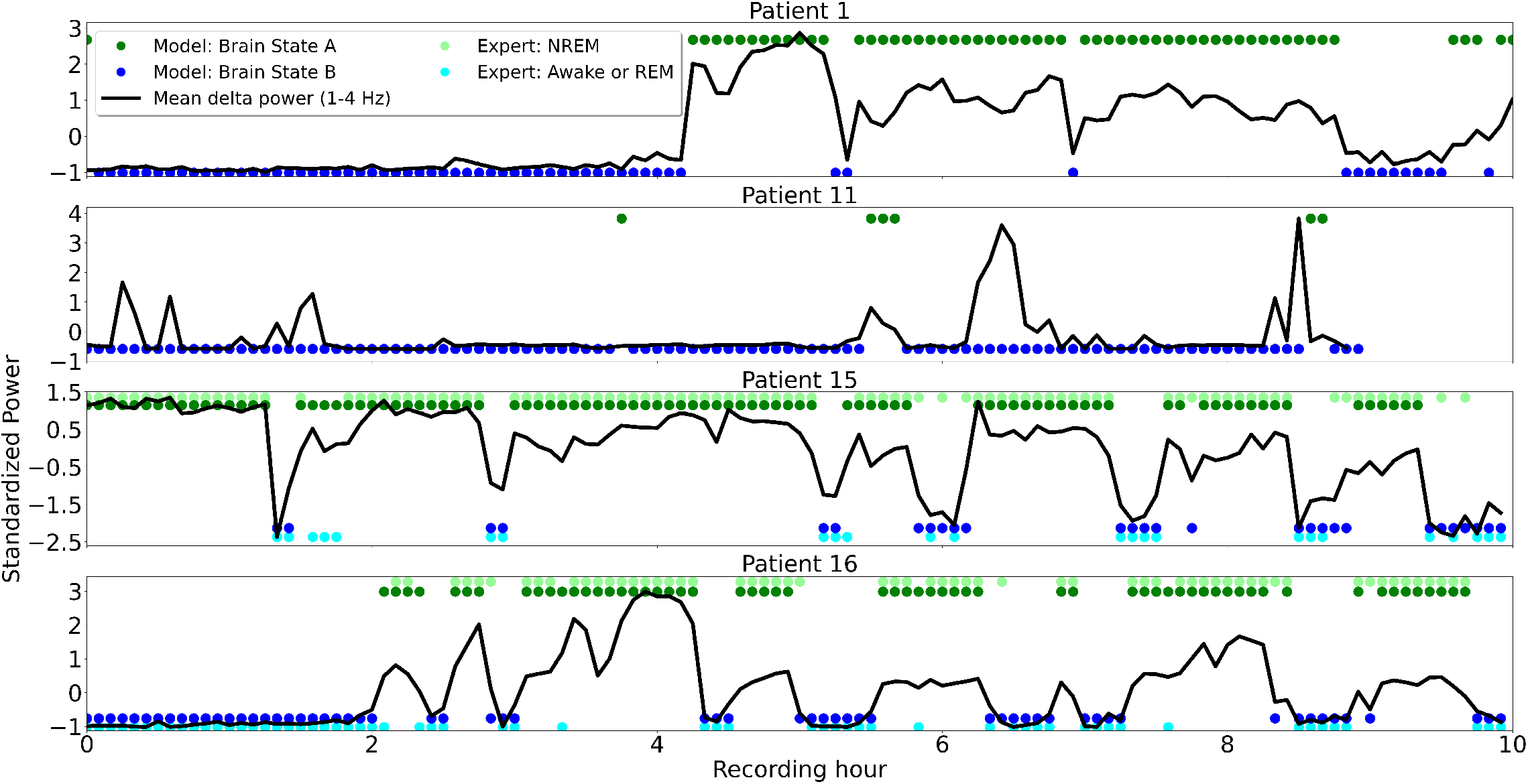
Model-derived brain states correspond to sleep and wakefulness. Representative examples are shown from Patients 1, 11, 15, and 16 with brain states A (dark green dots) and B (dark blue dots) obtained automatically every 5 minutes from Model 2. The labels of states A and B were assigned using the mean slow-wave delta power (1-4 Hz; standardized mean across channels), with brain state A containing higher delta power. Black lines represent the standardized mean delta power across channels. In patients 15 and 16, who had concurrent iEEG and scalp EEG, the HFO model-derived brain states differentiated slow wave sleep (i.e. NREM sleep stages 1, 2 and 3; denoted by light green dots in the upper portion of the bottom two subplots) from all other states (REM sleep and wakefulness; denoted by light blue dots in the lower portion of the bottom two subplots). This determination was made based on a comparison to expert sleep staging.

**Figure 8:**
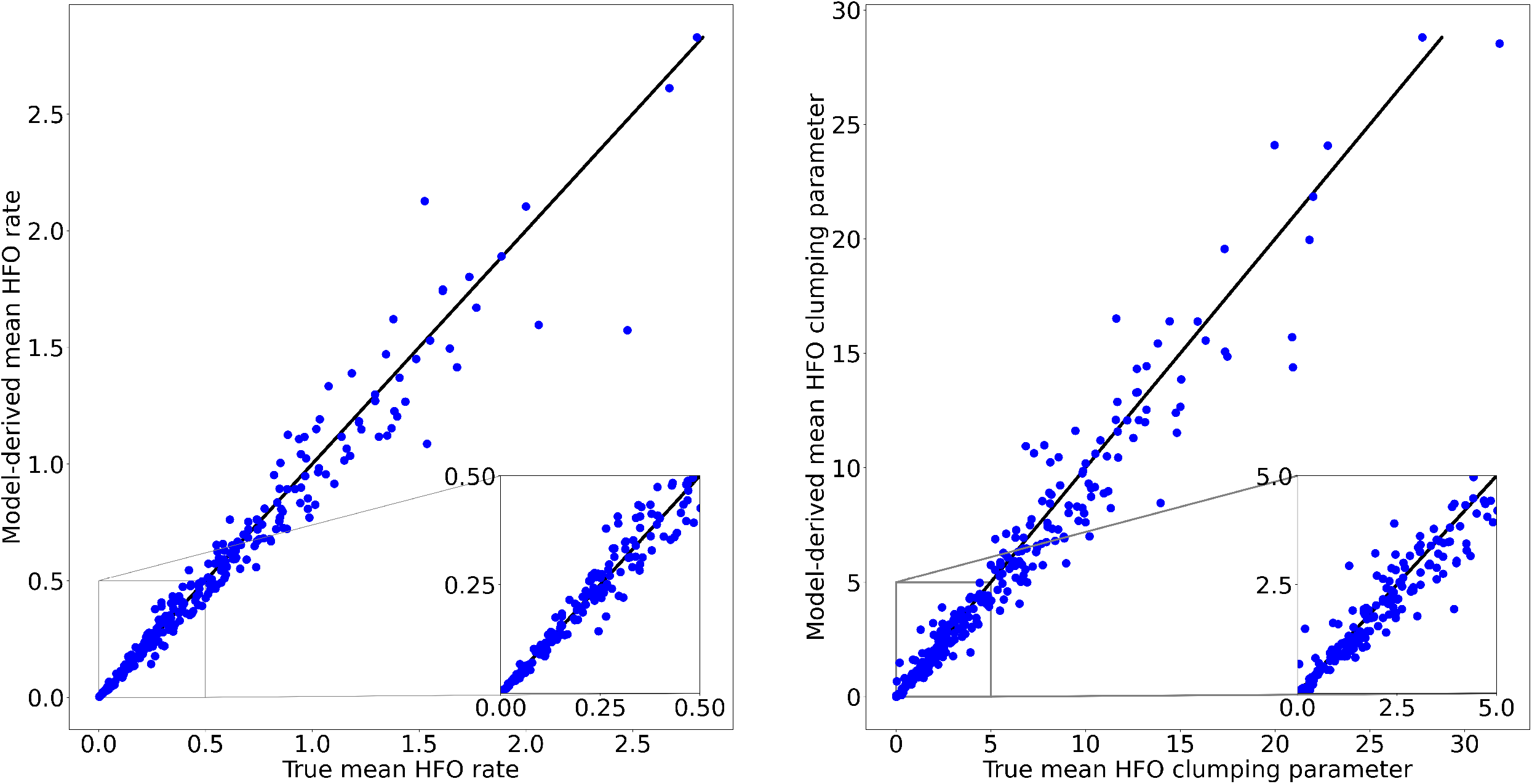
Recovery of mean HFO rates *λ* (Left) and mean HFO clumping coefficients (CC) *ζ* (Right) when simulating *E* = 300 channels over 5 hours of data from Model 1. The line corresponds to perfect parameter recovery. The insets zoom in on more common values.

**Figure 9:**
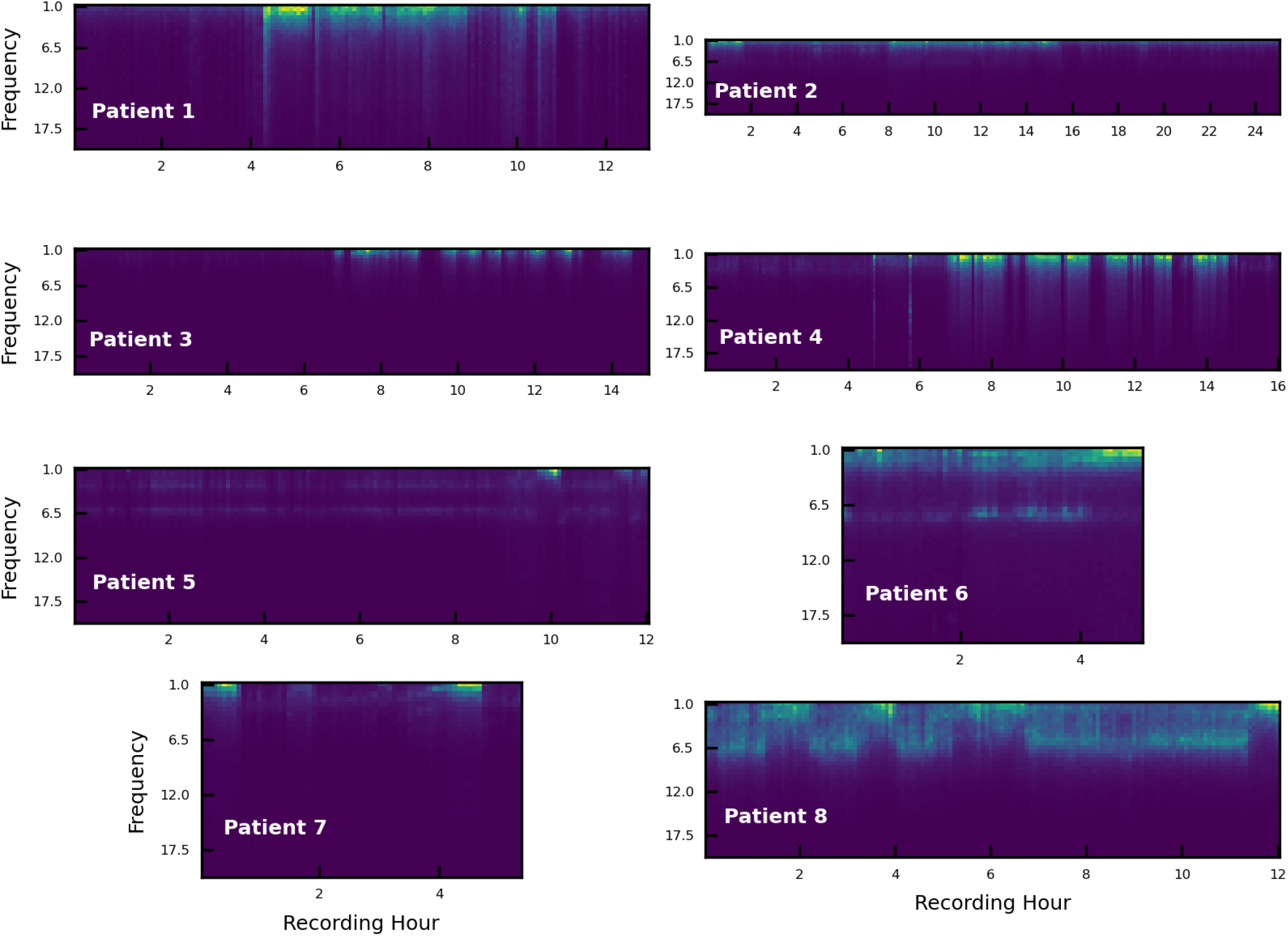
Relative power is shown from 1 to 20 Hz in a representative electrode within grey matter for Patients 1-8. The relative power was calculated for each 5 minute window over the duration of the entire recordings used in the analysis. Qualitatively, all patients had periods of increased delta power which suggests periods of sleep. However for all patients besides Patients 16 and 17, there was no ground-truth expert sleep staging to which to compare the relative delta power. See also **Figure 7** in the main paper.

**Figure 10:**
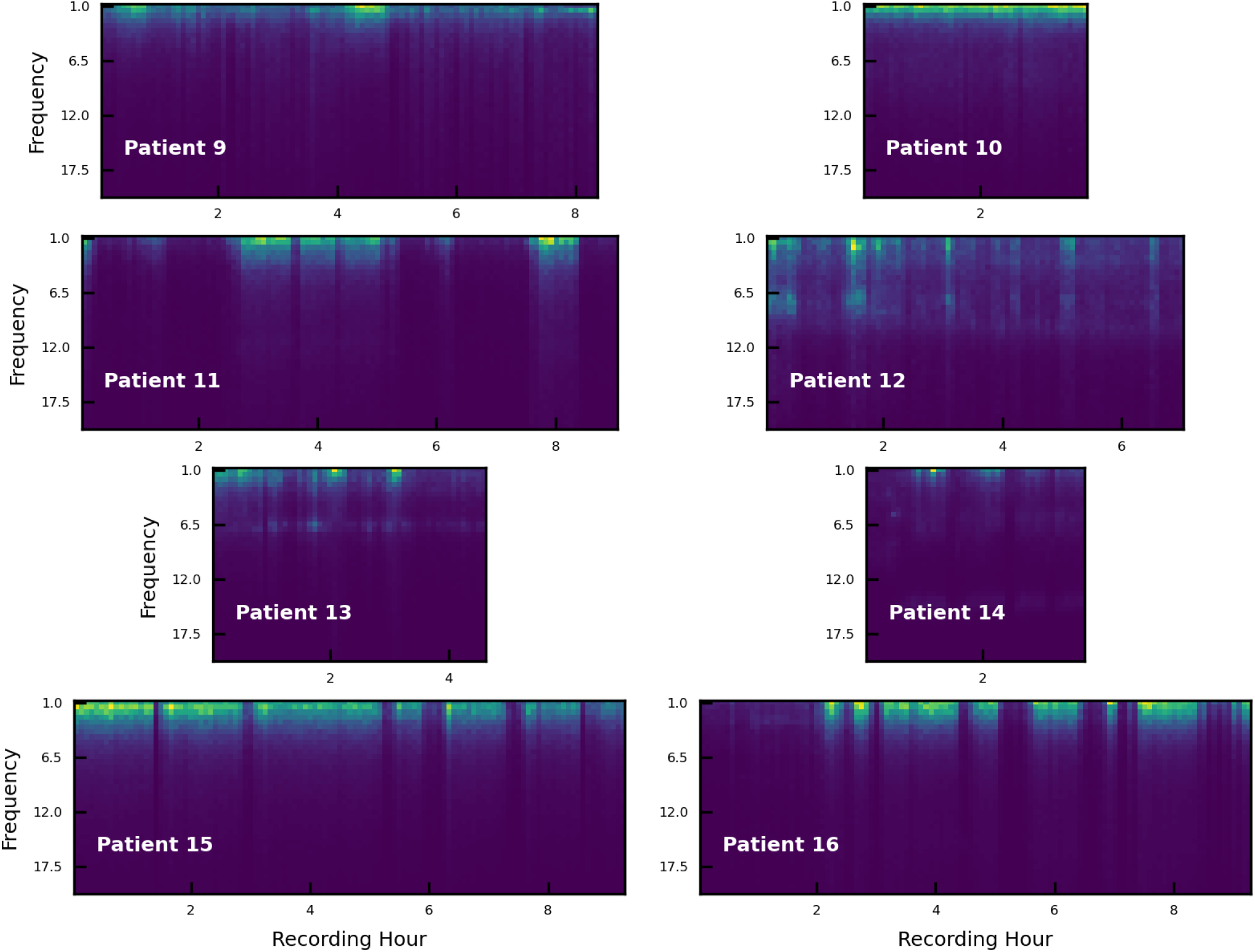
Relative power is shown from 1 to 20 Hz in a representative electrode within grey matter for Patients 9-16. The relative power was calculated for each 5 minute window over the duration of the entire recordings used in the analysis.

**Figure 11:**
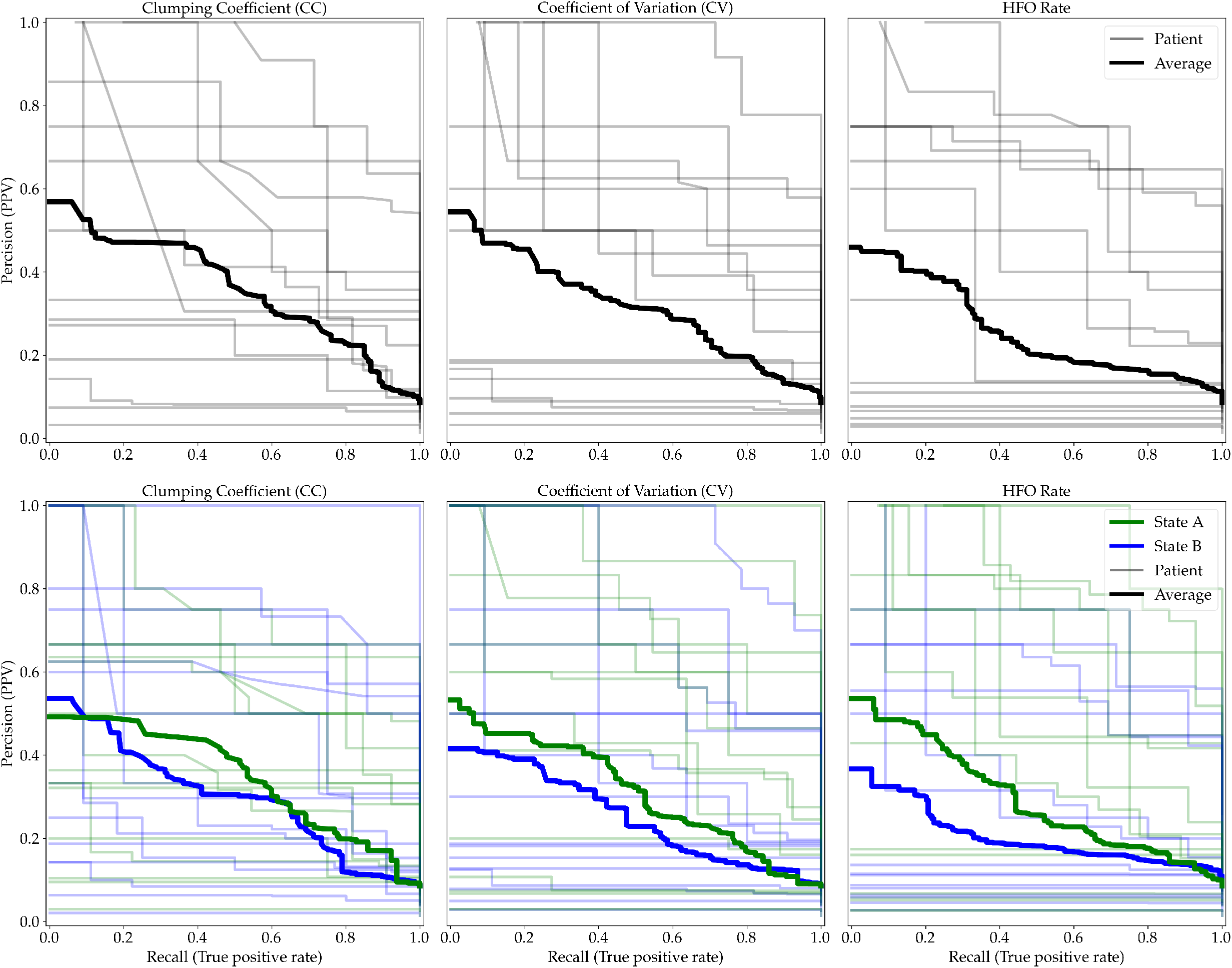
Recall (True Positive Rate) versus Precision (PPV) for each parameter. SOZ prediction was based on the clumping coefficients (CC; Left), coefficients of variation (CV; Middle), and HFO rate (Right) estimated by Model 1 (Top) and Model 2 (Bottom). ROC curves for individual patients (*N* = 16) are displayed using fine lines, and the average is shown in bold. From Model 2, the brain state with the most delta (1-4 Hz) power in each patient was labeled State A (green lines), while the other model-found brain state was labeled State B (blue lines).

As previously mentioned, we labeled brain state A as the brain state that contained the largest mean delta (1-4 Hz) power. However, we have reason to suspect that in at least one patient this automatic labeling failed and placed the majority of NREM sleep in brain state B. For instance, the HFO count data used in the models from Patient 11 was derived from 9 hours of neural recordings starting at approximately 22:45 at night, suggesting that the majority of the HFO count data should be from NREM periods. However, the largest delta power was contained in the brain state that occurred infrequently (see second panel of **Figure 7**). Note that this patient did have a significant Kruskal-Wallis test (*p* = .025) using a cutoff of *α* = .05, suggesting a difference in delta power between the two states. However this mislabeling might explain why Patient 11’s HFO parameters during brain state B were more predictive than brain state A, with the largest difference seen in the AUCs based on the CC (see **Figure 3**).

### 5.5. Assuming two brain states improved SOZ prediction for some patients

We found that assuming two brain states in hierarchical Negative Binomial mixture-models improved SOZ prediction in some patients by isolating NREM sleep automatically. This can been seen when comparing the SOZ prediction using parameters of Model 1 to Model 2 after fitting these models to data from each patient. For instance, the clumping coefficients (CC) of Patients 5 and 7 from brain state A in Model 2 result in higher AUC values than the CC of Model 1 (see **Figure 3**). This supports prior findings that NREM sleep should be used for not only calculating pathological HFO rates (Dümpelmann et al., 2015; von Ellenrieder et al., 2016, 2017), but also for calculating pathological clumping coefficients.

However for patients *overall*, there was not a clear benefit of using Model 2 over Model 1 because the data across patients was generally predictive of SOZ. We compared the values of FPR at which the TPR was 1, and we did not find evidence for a mean difference derived from CC1 versus CC2A (*p* = .337, *BF* = 0.39, two-sided paired samples t-test) nor CV1 versus CV2A (*p* = .993, *BF* = 0.26). However, we did find some evidence for a mean difference in the FPRs derived from CC1 versus CC2B (*p* = .017, *BF* = 3.55) and CV1 versus CV2B (*p* = .004, *BF* = 10.80). Similarly, we did not find evidence for a mean difference in AUCs derived from CC1 versus CC2A (*p* = .724, *BF* = 0.27) nor CV1 versus CV2A (*p* = .822, *BF* = 0.26), while we did find evidence for a mean difference AUCs derived from CC1 versus CC2B (*p* = .006, *BF* = 8.73) and CV1 versus CV2B (*p* = .010, *BF* = 5.45). We observed no significant mean differences between the SOZ prediction results using HFO rate based on Model 1 versus brain state A in Model 2 nor Model 1 versus brain state B in Model 2.

A similarity of prediction between the two brain states in Model 2 could be explained by consistent relative differences between channels. For example, although it is known that the rate of HFOs increases during NREM sleep (von Ellenrieder et al., 2017), the classification accuracy based on the two brain states could be similar if the relative rates between channels remain the same. To test this, we first compared the means of the three derived parameters across the two brain states in Model 2, collapsed across patients and channels, and we found that they were all significantly different. The CC had a mean and standard deviation of 3.48 ± 7.62 in brain state A and 5.24 ± 8.85 in brain state B collapsed across patients and channels (*p <* .001, *BF* ≈ 1.337 * 10^12^, two-sided paired samples t-test). The CV had a mean and standard deviation of 2.58 ± 2.70 in brain state A and 2.97 ± 1.63 in brain state B collapsed across patients and channels (*p <* .001, *BF* ≈ 6.583 * 10^6^, two-sided paired samples t-test). Note that the mean CC and CV values are quite larger than the smaller predictive values (see **Figure 4**) because SOZ channels made up only a small percentage of total channels in our study. The HFO rates had a mean and standard deviation of 0.59 ± 0.56 per second in brain state A and 0.47 ± 0.54 per second in brain state B, collapsed across patients and channels (*p <* .001, *BF* ≈ 4.903 * 10^8^, two-sided paired samples t-test). Then, to test whether the model-derived brain states captured independent information about HFOs, we calculated the Pearson correlation between the two brain states for the model-derived values of CC, CV, and HFO rates. The mean correlation coefficients of these measures indicated that HFO dynamics were similar across the two brain states in most patients, although there was a large range of correlation values. The Pearson correlations between brain states for all patients were as follows: *ρ*_*ζ*_ = 0.50 ± 0.25 (mean ± standard deviation) for CC, *ρ*_*γ*_ = 0.69 ± 0.24 for CV, and *ρ*_*λ*_ = 0.63 ± 0.28 for HFO rates.

## 6. Discussion

### 6.1 HFO clumping is a more reliable predictor than HFO rate

HFOs that occur with *less clumping behavior* (i.e. HFO occurrences that more closely follow a Poisson process) are more consistently predictive of SOZ than a high rate of HFOs.

The clumping coefficient (CC) and coefficient of variation (CV) were measured using hierarchical Negative Binomial models. Small CC and small CV were predictive of SOZ in most patients. In a second model, we found two CC and CV per iEEG channel using a model of two brain states obtained from a mixture of Negative Binomial distributions of HFO counts. CC were found to be more predictive of SOZ in the brain state corresponding to large delta power, likely corresponding to NREM sleep in at least half of the patients. Although CC based on all the interictal data using Model 1 were also predictive of SOZ. High HFO rates were also informative of the SOZ, but were less consistently predictive across patients than CC and CV. Our results also suggest that if HFO CC, CV, and rates from a single brain state are to be used in prediction of the SOZ, they should be assessed during NREM sleep. This supports previous findings in the field (Dümpelmann et al., 2015; von Ellenrieder et al., 2016, 2017).

### 6.2 Towards automatic classification of SOZ with interictal HFOs

Originally, we hypothesized that using mixture-modeling to automatically identify periods of NREM sleep would produce better prediction of SOZs. However, we did not find evidence that mixture-modeling greatly improved SOZ prediction, compared to Model 1, over all patients. While there were other differences between these two models, the benefit of mixture-modeling for clinical evaluation is not clear, compared to calculating parameters such as the clumping coefficient (CC) using all available data. These findings could be conflated if some patients had either only periods of wakefulness data or NREM sleep in the interictal subsets of data used in the modeling. This could be one reason why the delta power (1-4 Hz) of only half of the patients was significantly different between the two brain states. Qualitatively, all patients had periods of increased delta power (see Supplementary Figures 2 and 3). However, relative delta power could only be compared to ground truth expert sleep staging in Patients 16 and 17. Other recent studies have used quantitative methods to sleep stage iEEG data and compared the results to expert sleep staging in all subjects (Reed et al., 2017; Kremen et al., 2019).

On the other hand, in some studies it may be desirable to identify periods of NREM sleep. In these patients, fitting mixture models is an effective way of obtaining information about HFO dynamics without the need for concurrent EEG and manual sleep staging. Using the techniques presented here, there was no need to sleep stage the data (such as in von Ellenrieder et al., 2017) because the Negative Binomial (and Poisson, see Supplementals) mixture models automatically identified changes in HFO dynamics over time. Approximate sleep stages were automatically obtained as a result of the distribution demixing. Our method of performing SOZ classification with HFO mixture modeling discussed in this paper should be compared to (1) differentiating NREM from REM and awake prior to HFO rate analysis, (2) analyzing HFO dynamics coincident with high delta (1-4 Hz) power as a proxy for NREM sleep, and (3) using automatic iEEG sleep-staging (Reed et al., 2017; Kremen et al., 2019). The similarity of HFO rates in REM sleep compared to HFO rates during wakefulness has previously been shown (Staba et al., 2004). In two patients, we found that the dynamics of HFOs during REM and wakefulness are often similar within each channel. And in at least half the patients, this model-derived brain state B, the brain state with the smaller delta power (1-4 Hz), likely reflects REM and wakefulness because there was a significant difference in delta power between the two states. And we confirmed that model-derived brain state A did reflect NREM sleep in two of these patients whose data was sleep staged. It is also possible that the mixture modeling captures interictal HFO dynamics *independent of sleep stage* that are predictive of SOZ. However, this possibility should be explored further in other datasets.

### 6.3 Limitations of this study

Differences across patients and channels (such as differences in electrode size and locations, differences in brain shape and volume conduction, differences in disease state, etc.) may all play a role in the potential of HFOs to predict pathological tissue. In our data, most electrodes were placed in the temporal or frontal lobes based on the expected locus of epilepsy and other surgical considerations, and thus our approach may need to be validated using data from other cortical locations. In addition, we did not show that the SOZ could be predicted in Patient 14 using any parameter (e.g. see **Figure 3**). This patient had only one channel identified as SOZ by clinicians, making it difficult to evaluate the prediction of SOZ and possibly led to these near-chance outcomes. However the methods presented in this paper were promising for the limited number of patients and brain regions explored in this study. The importance of validating the predictive nature of these methods in additional patients is obvious and cannot be overstated.

Note also that only 4 of the 16 patients were completely seizure-free after surgery, denoted by Engel Outcomes IA in **Table 1** (Engel, 1993). This is consistent with the general notion that treatment of the seizure onset zone is often insufficient to prevent the occurrence of seizures. Moreover, in some patients, the SOZ could not be completely removed during surgery. Therefore future studies should include a more detailed analysis of electrodes within the resected volume in order to make a quantitative comparison to surgical outcome. This is a more valuable test of clinical utility, as it evaluates the ability of the quantitative method to identify the epileptogenic zone, rather than the seizure onset zone (for which standard clinical criteria already exist).

The choice of automatic detection algorithm and detection parameters will also have a significant impact on the results. We chose a simple algorithm, due to the large amount of data to be analyzed, but implementation of a more complex algorithm with post-processing steps to reject false positive detections based on the time-frequency decomposition may improve the specificity of the detection and classification of SOZ channels. For example, if the HFOs occur in a regular, oscillatory pattern, the temporal dynamics may appear more random or clumped with the addition of false positives due to artifacts. The use of a more specific detector may also enable the application of these methods to scalp EEG (Zelmann et al., 2014; von Ellenrieder et al., 2014; Kobayashi et al., 2015; Gotman, 2018; McCrimmon et al., 2021), as false positive detections due to muscle artifacts would be a source of noise when assessing HFO dynamics on the scalp (Nunez et al., 2016; Bernardo et al., 2018).

In our analysis, we treated all detected events equally, without attempting to separate pathological and physiological HFOs (e.g. see Liu et al., 2018). It is possible that these two types of HFOs have similar rates, but different temporal dynamics, in which case our proposed method could help distinguish between them. However, here we could only classify events as being inside and outside the SOZ, which would include both physiological HFOs and artifacts. Therefore, this question must be more explicitly studied with cognitive paradigms to elicit physiological HFOs, or the analysis could focus on the fast ripple frequency band (250-500 Hz), which is hypothesized to contain only pathological HFOs.

There may also be differences in pathological HFO dynamics between intracranial depth electrodes and cortical surface electrodes. These two types of sensors record from different amounts of cortical depth and volume, and intrinsic differences in neural behavior between different spatial scales could exist (Nunez et al., 2019). We might even expect differences in neural behavior between iEEG electrodes of different diameters at similar locations within the same patient, due to these reasons (Nunez et al., 2019). Lastly, we would expect some depth iEEG electrodes to be contaminated by noise, as the most lateral channels are sometimes outside the brain. There have been conflicting reports on the effect of electrode size on the ability to measure HFOs (Worrell et al., 2008; Châtillon et al., 2013); in this study, we collapsed across all types of intracranial electrodes.

Finally, our results contrast with previous results, such as work by Sumsky and Santaniello (2018), who found that bursting patterns of HFOs are more likely to be present in the SOZ. Both studies assumed that 1-second windows of HFO counts were described by particular count processes. However other modeling assumptions do differ between the two studies, which could lead to contrasting results. We used Negative Binomial models to parameterize the count process, while Sumsky and Santaniello (2018) used a non-stationary Point Process Model. We also built full ROC curves for SOZ classification, which were not used in the previous work. We therefore found it difficult to directly compare the classification results of both works. Further study is needed to understand differences in model predictions tested against large amounts of data.

### 6.4 Future improvements to algorithmic implementation

Faster methods of fitting Poisson and Negative Binomial mixture models are necessary for these methods to be applied in a clinical setting. In this study, we wished to fit hierarchical models in order to understand the relationship between channels and patients. However, in future studies, simple algorithms to fit mixture models of Negative Binomial distributions and other distributions, such as presented by (Nagode, 2015), may be sufficient.

Some models presented here did not converge as judged by the Gelman-Rubin statistic, 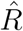,although the median posterior parameters were still informative for SOZ classification. This seemed to be due to the non-convergence of specific HFO rates and oscillatory dynamics for subsets of channels in some patients. This could be caused by artifacts being introduced into the HFO rates by the automatic detection process or due to actual physiological or pathological deviations from that channel’s rate in that brain state. It could be that the adaptive noise floor, which changed every 5 minutes within each channel using our automatic HFO detector (Charupanit and Lopour, 2017), injected artifactual HFO dynamics into the models. It is possible that fitting an HFO detector and Poisson / Negative Binomial hierarchical models *concurrently* would alleviate this convergence issue.

Model convergence is usually a bare minimum for hierarchical Bayesian model building. However, because the outcome of this study was SOZ classification and the non-converged models were still able to classify SOZ and non-SOZ, the results are still clinically relevant. Models that allow “noise” in the HFO dynamics to occur with some limited frequency could alleviate this issue. This could facilitate model convergence and may even yield better classification of the SOZ. In pilot analyses, we were unable to fit mixture models with three or four brain states in JAGS with sufficient convergence of chains. Thus, the resulting posterior distributions of HFO parameters were difficult to interpret. We are unsure if the data would be better described by a model with more brain states. Future work should seek to expand the number of brain states while allowing for artifactual HFO dynamics.

More complex hierarchical Bayesian models can be fit that provide further inference about the HFO dynamics and SOZ prediction. In particular, hierarchical Bayesian models that predict SOZ directly (instead of that prediction being derived from the posterior distributions of parameters) would be useful to assess the uncertainty in prediction. Also, it would be desirable to have a single hierarchical Bayesian model that includes all patients’ data, to understand commonalities across all patients and how SOZ prediction varies with individual differences in disease state. However, the computational load of this model would be particularly high with the multiple hours of HFO count data, and certain “big data” management schemes would have to be deployed. Future work should also seek to combine both HFO dynamics (CCs and CVs) and HFO rates for SOZ prediction within hierarchical Bayesian models, especially because some patients’ SOZ were better predicted by HFO rates (compare **Figures 3, 5, and 6**). This may suggest that complementary information for SOZ prediction is provided by HFO dynamics and HFO rates. Such hierarchical Bayesian models should be compared to similar endeavors to combine features for SOZ prediction during interictal periods using machine learning and artificial intelligence techniques (Varatharajah et al., 2018; Cimbalnik et al., 2019; Weiss et al., 2019). Finally, including other possibly predictive data such as delta (1-4 Hz) power, sleep stage, patient information, etc. directly into these hierarchical models could improve SOZ prediction. We felt as though many of these models were outside the scope of this paper, and each new model developed must be rigorously tested and tuned. Thus, we view this paper as the first step into a possible use of hierarchical Bayesian techniques in the prediction of SOZ with interictal iEEG data, and we look forward to further work on the topic.

## Acknowledgements

This work was supported by the NINDS of the National Institutes of Health under award number R01NS116273 as well as a grant from the American Epilepsy Society. The content is solely the responsibility of the authors and does not necessarily represent the official views of the National Institutes of Health. Casey Trevino is thanked for her help with implementing HFO artifact removal algorithm.

## 7. Supplementary Materials

### 7.1 SOZ prediction without qHFO algorithmic correction

We found that while prediction improved when using the qHFO algorithmic correction, most improvements were in patients without localization information. Thus we don’t think that the qHFO algorithmic correction is necessary for automatic prediction algorithms developed in the future. However further research with larger sample sizes of patients is necessary to understand the effect of the qHFO algorithms.

### 7.2 Negative Binomial process model recovery

The simulated mean HFO rates and mean HFO clumping parameters were recovered when simulating Negative Binomial processes when simulating *E* = 300 channels over 5 hours of data from Model 1 (see **Supplemental Figure 8**). However individual 5 minute HFO rates and HFO clumping parameters are not consistently recovered, and therefore caution should be taken when interpreting HFO rates and clumping parameters of small time segments. Note that individual 5 minute window parameters were not used for SOZ prediction, only the average of these parameters that recover the simulated mean parameters.

### 7.3 Poisson process mixture model

The exact model for each patient, which we will refer to as Model 3, is given by the following equations:

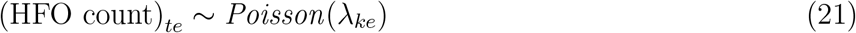

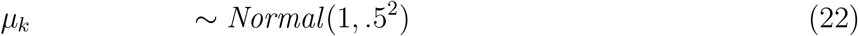

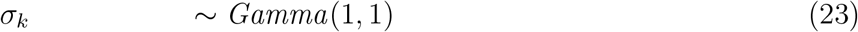

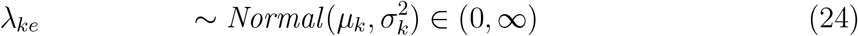

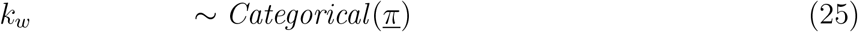

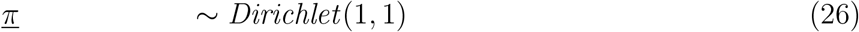

Such that HFO count per second *t* is described by a Poisson process with rate *λ* for each brain state *k* and channel *e*. Each rate *λ* is drawn from a hierarchical distribution with a mean rate *μ* for all channels in that brain state. The brain state *k* for every 5 minute window *w* is drawn from a Categorical distribution with a vector of probabilities *<u>π</u>* totalling 100% for all brain states (*<u>π</u>* is a vector of size *k* = 2 in the presented data for just two brain states).

Brain states and ROC curves obtained from from this hierarchical Poisson mixture-model were nearly identical to brain states and ROC curves obtained from Model 2 in early work. Therefore we do not report the results here. We reproduced the model in Supplementary materials as an easy comparison to the hierarchical mixture-model Negative Binomial Model 2.

### 7.4 Patient-specific results

### 7.5 Precision-Recall curves

Precision-Recall (PR) curves show the Positive Predictive Value (PPV; Precision) versus the True Positive Rate (Recall). PPV is defined as the number of true positives divided by the total number of true positives and false positives. The PR curves for each patient are provided in **Supplemental Figure 11**. PR curves have been suggested to be superior to ROC curves in evaluating binary classification because they are less influenced by unevenly distributed classes compared to ROC curves (Davis and Goadrich, 2006). In the data presented in this paper, the number of SOZ channels is much smaller than the number of non-SOZ channels, with the total fraction of SOZ channels being 112*/*1391 = 0.081 collapsed across patients. The large number of non-SOZ channels could inflate ROC curves while the PR curves could remain the same with fewer non-SOZ channels in the data.

In general, the results using PR curves mirrored the results based on the ROC curves, with CC and CV showing better prediction than HFO rate, and State A (corresponding to larger delta power) having better prediction than State B. However, PR curves cannot easily be compared across patients due to different baseline ratios of SOZ to non-SOZ channels. The minimum fraction of SOZ channels was 0.014 in Patient 14. The maximum fraction of SOZ channels was 0.26 in Patient 10. This implies that baseline chance prediction of the PR curves ranges from a horizontal line at 0.26 to a horizontal line at 0.014 in **Supplemental Figure 11**. We also had no reason to remove electrodes from the analysis to compare PR curves across patients, and thus we find these curves to be difficult to interpret compared to ROC curves. It is perhaps noteworthy that fewer PR curves are flat lines below 0.2 for CC and CV-based prediction compared to HFO rate-based prediction. Note that we did compare the performance of ROC curves based on real SOZ and non-SOZ labels to ROC curves based on randomly shuffled labels in the main body of the paper, with all summary ROC evaluation statistics given in **Table 2**.

